# Clusterdv, a simple density-based clustering method that is robust, general and automatic

**DOI:** 10.1101/224840

**Authors:** João C. Marques, Michael B. Orger

## Abstract

How to partition a data set into a set of distinct clusters is a ubiquitous and challenging problem. The fact that data varies widely in features such as cluster shape, cluster number, density distribution, background noise, outliers and degree of overlap, makes it difficult to find a single algorithm that can be broadly applied. One recent method, clusterdp, based on search of density peaks, can be applied successfully to cluster many kinds of data, but it is not fully automatic, and fails on some simple data distributions. We propose an alternative approach, clusterdv, which estimates density dips between points, and allows robust determination of cluster number and distribution across a wide range of data, without any manual parameter adjustment. We show that this method is able to solve a range of synthetic and experimental data sets, where the underlying structure is known, and identifies consistent and meaningful clusters in new behavioral data.

**Author summar:** It is common that natural phenomena produce groupings, or clusters, in data, that can reveal the underlying processes. However, the form of these clusters can vary arbitrarily, making it challenging to find a single algorithm that identifies their structure correctly, without prior knowledge of the number of groupings or their distribution. We describe a simple clustering algorithm that is fully automatic and is able to correctly identify the number and shape of groupings in data of many types. We expect this algorithm to be useful in finding unknown natural phenomena present in data from a wide range of scientific fields.

## Introduction

A notable feature in data is that the points, rather than being evenly distributed, are more densely clustered in some regions of space than others. These clusters may be distributed around single points, or more extended complex shapes. It is often necessary to determine how many clusters exist in a particular data set, and how points are distributed in these clusters, since this structure may reflect the natural processes underlying the data being collected. Unsupervised computational methods that can determine the number of clusters in data and define their natural boundaries are useful to identify unsuspected natural phenomena and are widely used across many disciplines of science.

Although clustering analysis has a long history, there is no universal consensus on the definition of a cluster or on which clustering algorithm is the most effective [1]. In fact, “an impossibility theorem for clustering” was formally proven, showing that there is no single clustering function that can satisfy three fundamental criteria, scale-invariance, richness and consistency [2]. In spite of that, it is possible to construct clustering algorithms that have broad applicability by relaxing the proposed criteria. One of the aims of machine learning is to develop general purpose clustering heuristics that function automatically for the most diverse types of data possible and hence many clustering strategies have been proposed [3,4].

Whereas some widely used clustering approaches depend on assumptions about the cluster shape, some density-based clustering methods allow clusters of arbitrary shape to be discovered [5]. One clustering method, based on search for density peaks (named clusterdp or DensityClust), is fast, resilient to noise, captures clusters of arbitrary shapes, and provides an intuitive method to select the number of clusters [6]. This method successfully solved artificial data sets contaminated with noise, and clusters with fuzzy edges and varying shapes. Also, it performed well in classifying faces from images and, in a meta-study of clustering methods, to solve real world biomedical data [4].

However, in many cases, the output of clusterdp is critically dependent on the parameter that is used for estimating the local densities (dc). Additionally, as we also show here, the clusterdp heuristic fails when applied to certain distributions with clearly distinguishable clusters. We designed an alternative approach that is both more general and allows cluster selection to be fully automated. Our method, which we call clustering by density valleys (clusterdv), is based on similar principles to clusterdp, but differs in several key elements. We use an adaptive Gaussian density estimator to compute the local densities, in common with other variants of the method[7], and we define point separation based on how deep a density valley had to be traversed to connect pairs of points. We developed a robust rule to identify, and to hierarchically order, putative cluster centers. Lastly, we implemented methods, based on statistical comparison with reference distributions and the largest jump within the cluster hierarchy, to select the number of clusters automatically.

We validated clusterdv by applying it, without parameter tuning, to a wide variety of artificial and real-world test data with known ground truth cluster identity. We show that clusterdv can identify the correct number of clusters very reliably, including in distributions that cannot be clustered using clusterdp. The method also assigns points to the correct clusters with high accuracy. Finally, we show that it allows robust identification of behavioral categories in experimental data from larval zebrafish. Altogether clusterdv is an automatic unsupervised method for density cluster identification that achieves state of the art performance over a wide range of types of data, making it an ideal tool to discover structure in real world data where the ground truth is not known.

## Results

### Identifying limitations of density peak clustering

The clusterdp algorithm relies in calculating two quantities: the local density (ρ) at each point and the minimum distance from each point to a point with higher ρ (δ) [6]. Both of these values depend on the choice of dc, a free parameter in the clustering method which determines the spatial scale used to calculate local densities. Since clusterdp requires the user to select a number of cluster centers based on the distributions of ρ and δ, we developed an automated heuristic for this choice that we named automatic clusterdp (see Materials and Methods and Fig. S1). In most cases, this method, when applied to data sets for which the ground truth was available, performed as well or better than human observers in selecting the correct number of clusters (Fig S2).

We took advantage of the good performance of the automatic clusterdp to sample the dc parameter for 8 data sets used in the original study and found that the results of clusterdp are highly dependent on dc, for all data sets we tested, and there was no single value or range of values that consistently gave correct results (Fig. S3).

Another potential pitfall of clusterdp is that, if δ uses Euclidean distance as its metric, it will tend to favor density peaks that are far apart within large dense regions of points over nearby peaks that are clearly separated by an empty region of space. Arguably, though, the latter is a more salient feature of the data. To illustrate this point, we constructed a synthetic data set, hereinafter called “exclamation mark 1”, that consists of groups of points drawn from two spatially uniform, rectangular distributions: an isolated group of low density points, very close to an extended high-density region (Fig. 1*A*). This situation is commonly observed in real data sets, for example the zebrafish swimming data described later, where groups may be very unevenly represented and the more spatially restricted cluster has much lower density. The clusterdp algorithm is unable to find the low-density cluster without also splitting the single dense cluster into several parts (Fig. 1B). This is due to the fact that clusterdp implicitly ranks cluster centers according to the values of δ and local density, making it impossible, in case of the “exclamation mark 1” example, for a user or an automatic algorithm to select the low density cluster center (red dot) without including first the two cluster centers that exist in the high density cluster (cyan and blue dots) (Fig. 1B). This problem is exacerbated by the fact that the originally proposed method for assigning points to clusters does not correctly partition the data, even when the cluster centers correctly identify both groups (Fig. 1*B*, third panel, dark blue points in the corner of the lower cluster). This failure to rank the cluster centers correctly does not occur in a data set with similar characteristics, but in which the smaller cluster has higher density (Fig. 1*C-D*). To verify that this limitation was not dependent on parameter choice, we systematically varied dc and plotted the locations of cluster centers for all cases where the correct number (2) was found. In all such cases, for the “exclamation mark 1” data set, the cluster centers were both found in the larger cluster (Fig 1*E*). The opposite result was observed for the “exclamation mark 2” data set where, if the algorithm found the correct number of clusters, they always spanned the two regions (Fig. 1*F*). These results suggest two key limitations of clusterdp: a sensitive dependence on parameter choice, and a failure to capture correctly the structure in certain simple data distributions.

**Fig. 1.**
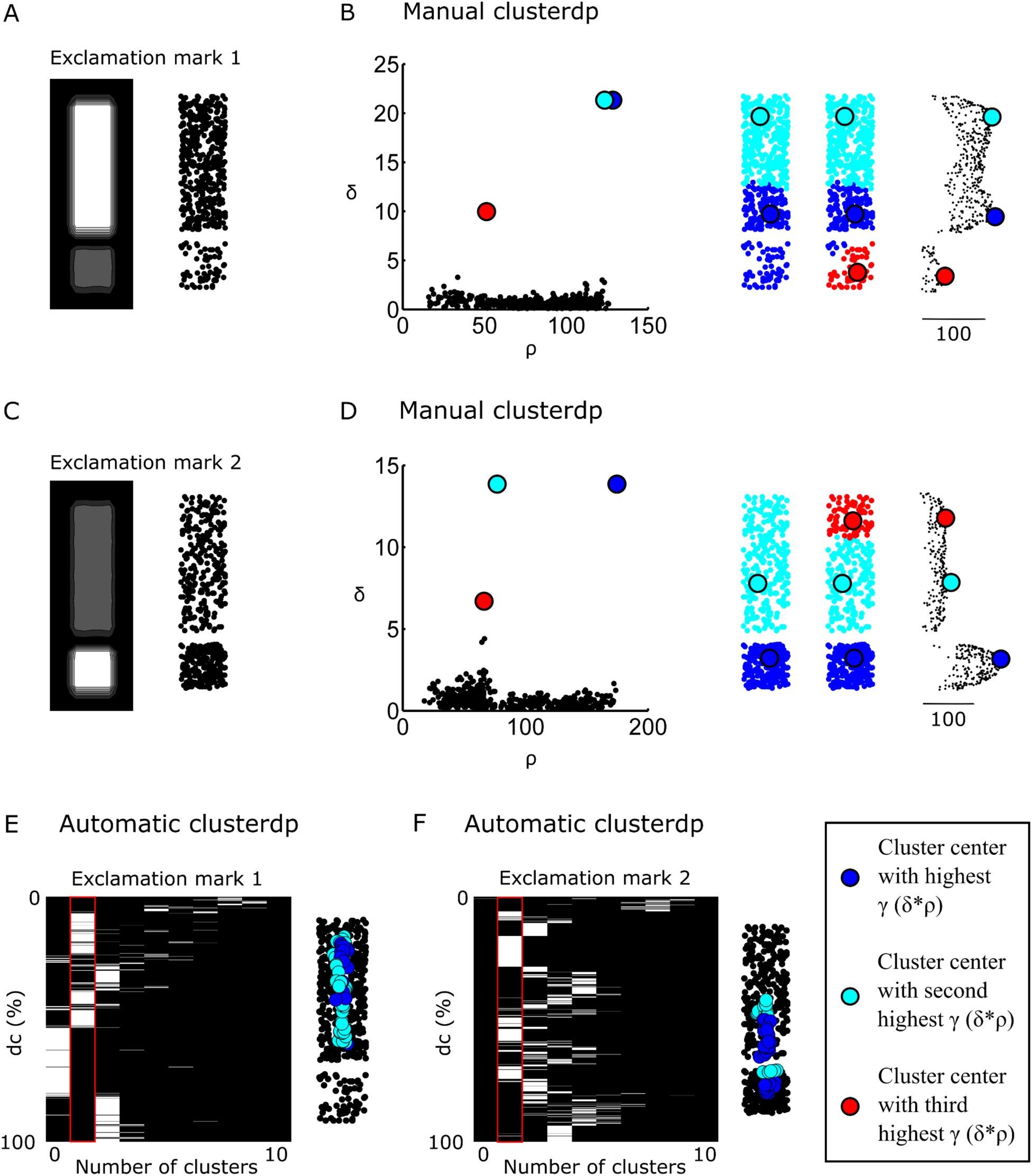
Density peak clustering fails with uneven clusters. (*A, C*) The “exclamation mark 1 and 2” data sets were drawn from two-part probability distributions (left). White represents 2.5-fold higher probability than grey and black is probability 0. (*B, D*) Manual clusterdp applied to the “exclamation mark” data sets. Left to right: clusterdp decision plot (ρ vs δ)) of the distribution in (*A, C*). Clusterdp solutions of data in (*A, C*) by picking the two or three cluster centers with highest γ (δ*ρ). Density profile of data in (*A, C*) (d_c_ = 9%). (*E-F*) Left: Number of cluster centers picked by automatic clusterdp in function of d_c_ value for the “exclamation mark 1” (*E*) and the “exclamation mark 2” (F) data sets. Red outlines mark the ground truth (2 clusters). Right: the cluster centers obtained by automatic clusterdp whenever the two-cluster solution was selected. Cluster centers and points are color coded blue-cyan-red in order of decreasing γ (δ*ρ) as in legend.

### The clusterdv algorithm

We set out to create a new clustering heuristic, based on the working principle of clusterdp, that was robust, general and automatic. That is to say, it should give repeatable results across many different data distributions, without the need for parameter fitting and should select the most suitable numbers of clusters in which to partition the data without human intervention. First, to calculate the local density at each point we used an adaptive Gaussian-based kernel density estimate [8], with the bandwidth at each point based on a heuristic that was applied identically, without any parameter tuning, to all data sets in this report (Fig. 2*A*-*B*). Similar adaptive approaches have been used in other clusterdp adaptations[7]. Secondly, we defined the separation of pairs of points in a way that aimed to capture a notion of distinct clusters. Specifically, two points should be considered well separated if you have to pass through a region of low density to get from one to the other, regardless of how close together they are in space. This separation was quantified by finding the path connecting two points, within a graph spanning the whole data set, along which the minimum density was highest. We first estimated the minimum density along lines defined by single edges in the graph by sampling at discrete intervals from the previous kernel density estimate (Fig. 2*C*). After, we searched for the path joining each pair of points, following any of these lines, which dropped the least in density along its length (density paths, Fig 2D). To find such paths we used the single link algorithm [9] on the previously calculated density lines. Therefore the value of the ‘density path’ between two points is the lowest density you have to pass through to get from one point to the other. Since it should be possible to join two points that lie in the same unimodal cluster by a path whose density profile is always higher than the lower density point, a point from which you cannot get to a higher density region, without passing through a region of lower density first, is at a local maximum, and therefore should be considered as a potential cluster center. So, for each point, we calculated the highest density path value between that point and any point of higher density (maximum density valley depth) (Fig. 2*D*). Figure 2*E* shows this maximum density valley depth, plotted against the local density at each point. However, in order to be sensitive to clusters of very different densities, we should not consider the absolute value of the density drop, but rather how low it is relative to the associated peak. We chose to consider a low-density peak, separated by an empty region from the rest of the data, as a more salient feature of a data set than a small dip separating two very high-density peaks. To capture this distinction, we calculated a separability index (SI) by first dividing the maximum density valley depth of each point by its local density, in effect normalizing the drop of density to the density at each point, and then subtracting that value from 1 (see materials and methods for formula). A SI value of 1 indicates that there is a region of zero density between a point and any higher density region, and a negative value indicates that a point has a path to a denser point that never dips below the starting density and is therefore not a cluster center (Figure 2F). All putative cluster centers (points with positive SI) can now be ranked according to their SI. These points are arranged in a hierarchical tree by connecting each new center point with the branch it would be assigned to, if the clusters were assigned without using that point (Fig. 2*G*). This SI tree reflects the hierarchical organization of clusters that exist in data. For data sets with complex clustering structure, for example with nested groups of clusters (Fig. S4), the SI tree will capture this organization through groups of nodes at different levels. The total number of clusters is determined by distinguishing from the pool of “putative” cluster centers, which ones are “real” and which arise due to stochastic variations in the density estimation of data with finite sample size. In practice, the selection of “real” clusters is achieved by setting a cut-off on the SI values (blue line in Fig. 2*F-G*). We applied clusterdv to the “exclamation mark” data sets and confirmed that the SI value ranks cluster centers correctly for both data sets (Fig. 2*I*-*J*). Additionally, if more than two cluster centers are selected (red lines in Fig, 2I-J), the data is partitioned in a manner than respects the true cluster boundaries (left panels of Fig. *I-J*), unlike clusterdp (Fig. 1). In summary, clusterdv does not need any parameter optimization to work and reports the ranking of cluster centers correctly, even in difficult cases such as the “exclamation mark 1” data set.

**Fig. 2.**
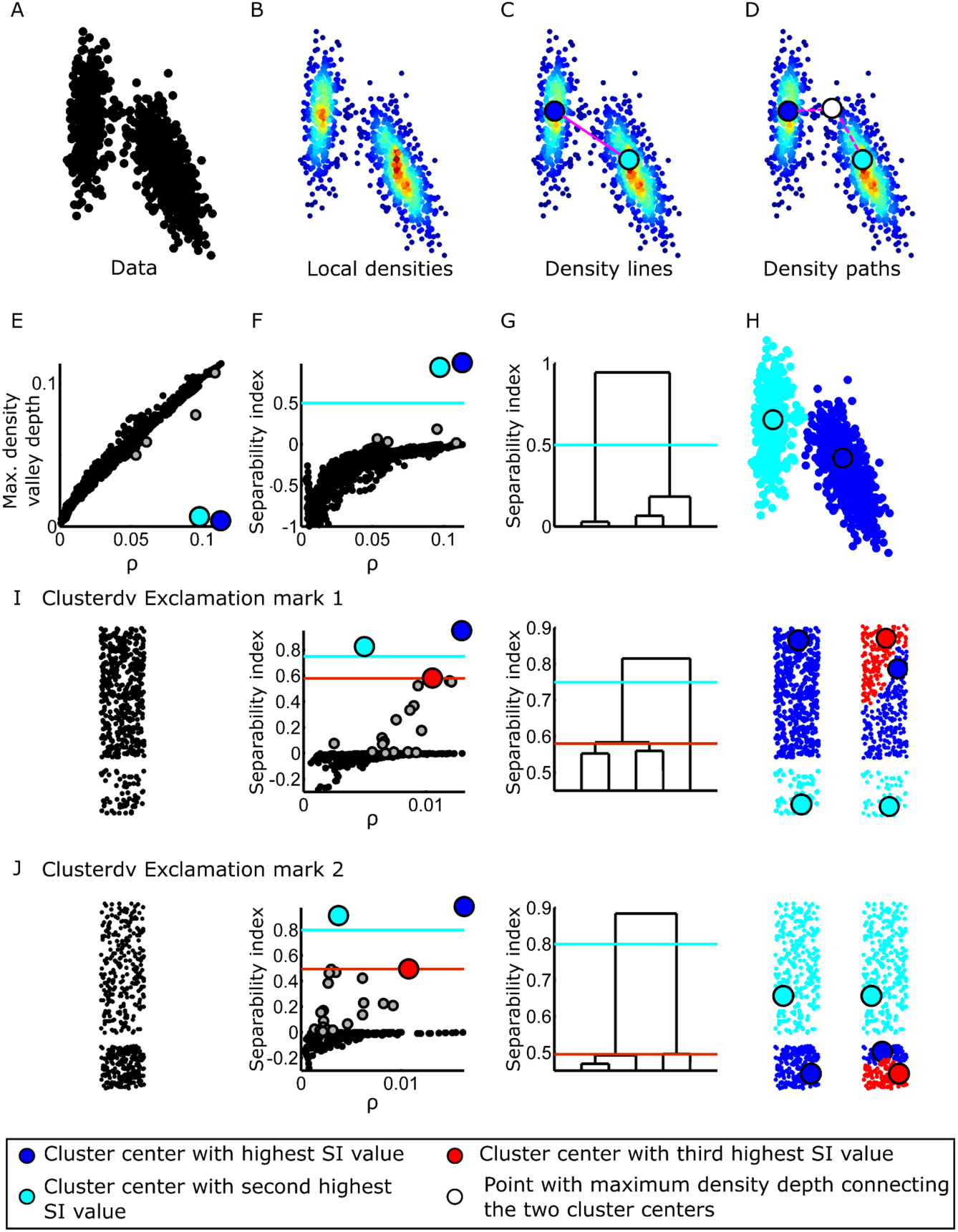
The clusterdv method. (*A*) Point distribution drawn from a mixture of two gaussians. (*B*) Local densities are calculated using an adaptive gaussian density estimator. (*C*) Density profiles are calculated, in a set of discrete steps, along straight lines between pairs of points. (*D*) The single link algorithm is used to determine the density path value, the highest minimum density along a path that connects one point to another, via this set of straight lines (path shown by white point). For each point, the maximum density valley depth is the highest such value connecting that point to a point of higher density. (*E*) Maximum density valley depth versus local density (ρ). (*F*) Separability index (SI) versus local density (ρ). Paths that don’t have a dip in density can give negative values because the two points are in the same cluster and the end points are not considered. Blue line is a manually selected cut-off that selects the “real” cluster centers (cyan and clue circles). Grey circles are “sporadic” cluster centers that were not selected. (*G*) Dendrogram computed from separability index. (*H*) Cluster assignment of the point distribution in (*A*) obtained by choosing the cluster centers with higher SI value than the blue line in (F-*G*). (*I*-*J*) Left to right: distribution, clusterdv decision plot (SI vs. ρ), SI dendrogram, and two and three cluster solutions for the exclamation mark 1 (*I*) or 2 (*J*) data sets. Cluster centers and points are color coded blue-cyan-red in order of decreasing SI value as shown in legend.

### Validation of clusterdv

Clusterdv is able to solve data sets with uneven clusters where the tightest cluster has lower density, but can the correct SI cut-off be determined automatically? and does the method give correct solutions for data with other characteristics? To answer these questions, we applied clusterdv to 33 artificial and real-world data sets with known ground truth that were designed to present varying difficult challenges for clustering analysis (Table S1). Three automatic criteria were tested to determine at which level the SI dendrogram should be cut and decide the number of clusters in each set. The more conservative method was to choose the largest jump in SI in the cluster tree. Alternatively, we estimated the SI distribution of clusters identified by chance in control data sets with similar spatial extent (“simplex”) or density profiles (“onion”), but with only a single density peak in the underlying distribution (Fig. S5 and Fig. S6) and selected all clusters above these thresholds (see Materials and Methods for details). The three methods gave the same correct solution for 13 data sets, and, with the exception of the Olivetti face data set [10], spanned a range of cluster numbers encompassing the known correct value. The max SI jump, in every case, gave a higher SI cut-off than the other criteria and was able to solve correctly 29 of the 33 data sets (Fig. 3). There were only four data sets for which the SI jump criteria failed to find the exact solution, and in three of those it identified a number of clusters just one step away from the correct answer. It is important to note that, since several of the data sets are real world data sets, it is not clear either that the labeled groups are the only clusters that should exist in the data, or that there should exist a clear distinction between clustered and non-clustered organization, which the SI jump assumes (Table S1). The methods based on reference distributions usually identified more structure, especially in the real-world data sets, as might be expected, but still also had high success rates (Table S1). In the case of the Olivetti face data set [10], the clusters were fractured in the feature space that was used [11], so it may not be possible to improve on this result without a different representation of the data. Nevertheless clusterdv achieved higher success (Olivetti, SI jump, FMI = 0.77 and Fig. S7), with much larger true associations for a given false positive rate, than previously published methods [6] (78 percent at a false alarm rate of 1, versus approximately 65 percent previously reported for clusterdp). Another notable feature of these results is that the correct solution almost always existed within the dendrogram of cluster centers, and in 31 out of 33 cases could be obtained by cutting at a single level (shown in best cut-off column of Table S1). In the case of the MNIST handwritten number data set [12], the ten digit solution existed in the tree, but could not be found using a single SI cut-off, without also finding multiple clusters within digit groups. The ‘correct’ solution could however be found by choosing the most balanced partition into ten clusters, and this could be used as a classifier which gave a 5.9% error rate for new data. This performance is comparable with the best unsupervised methods described for MNIST, that also use information about the construction of the data set, but not individual data labels (5% error rate in [13]) (Fig. S8). Importantly, in all cases for which the correct number of clusters was identified, the assignment of cluster centers always matched the ground truth, and the assignment of points to clusters was largely correct (Table S1).

**Fig. 3.**
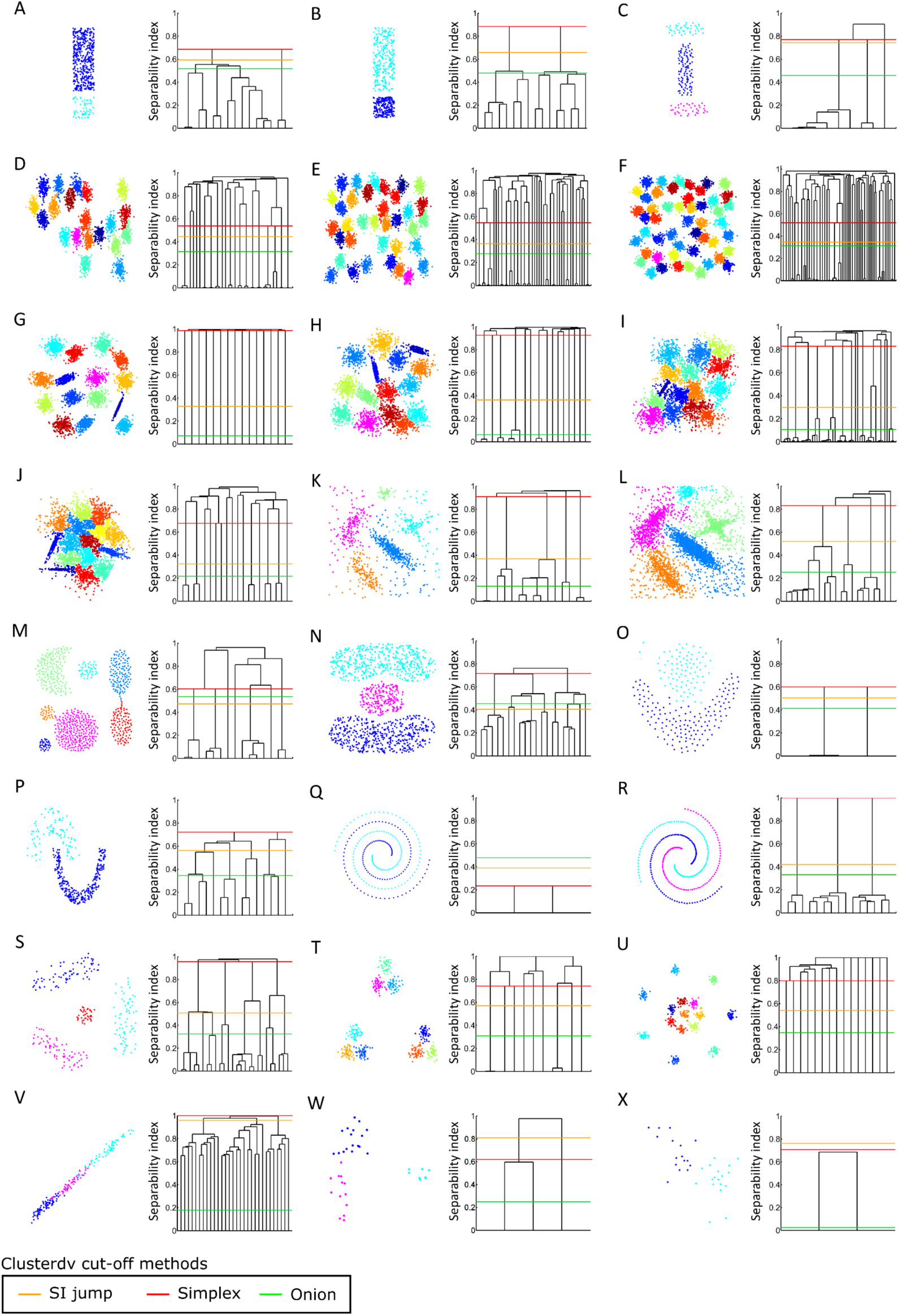
Clusterdv gives the correct solution for artificial and real-world data sets. (*A*-*U*) Synthetic data sets: (*A*-*C, T*) data sets from this study, (*D*-*F*) [20], (*G*-*J*) [23], (*K*-*L*) [6], (*M*) [16], (*N*) [4], (*O*) [17], (*P*) [18], (*Q*) [4], (*R*) [19], (*S*) [4], (*U*) [21]. (*V*-*X*) Real-world data sets: (*V*) seeds [24], (*W*) bone marrow [22], (*X*) Zachary [25]. Left panels: cluster assignment of the data sets according to the SI jump criteria. Right panels: SI dendrograms. Lines show cut-off criteria according to legend.

### Clusterdv outperforms clusterdp

Although clusterdv is capable of automatically, without any parameter tuning, solving a wide range of data sets, how does its performance compare with clusterdp, the state of the art density-based clustering method? To test this, we applied clusterdp, under the recommended range for the dc parameter [6], to the same array of data sets previously used to test clusterdv. We started by testing the automatic clusterdp against the three automatic methods of clusterdv. Clusterdv using the SI jump criteria outperformed automatic clusterdp, and also the two reference-based cut-offs (Fig.4A), achieving significantly higher values of FMI than any other method. Since the automatic rules to choose the number of clusters vary between clusterdp and clusterdv, we performed an additional test where the number of clusters is set, for each clustering method and data set, according to the ground truth. In this case, clusterdp and clusterdv were given the correct number of clusters (best cut-off). In this case, any problems in solving correctly the data sets are not due to the heuristic that chooses the cluster number but must come from problems in other steps of the algorithms. Fig. 4B shows that, given the best possible cut-off to choose the cluster number, clusterdv still clearly surpasses clusterdp, being able to correctly solve 31 of the 33 data sets, while clusterdp could only solve correctly 21 (Table S1). Overall, clusterdv managed to outperform clusterdp in one of its automatic implementations (SI jump criteria), but also when the number of clusters was optimal, so the gain of performance for clusterdv likely comes from having a better method to estimate the local densities and having a more robust rule to rank the cluster centers than clusterdp.

**Fig. 4.**
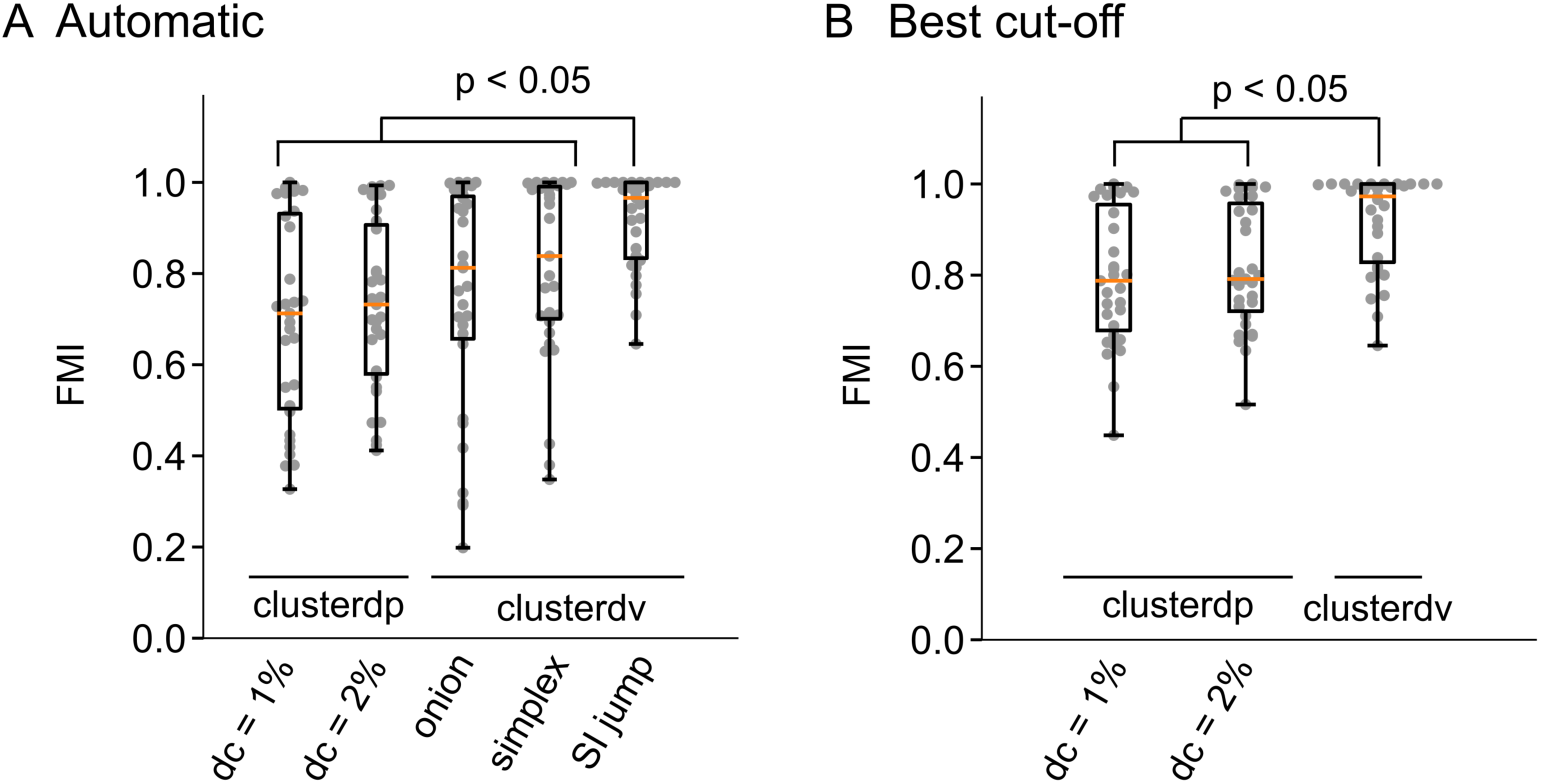
Comparison between clusterdp and clusterdv. The clusterdp and clusterdv methods (legends in x axis) were applied to 31 data sets that had known ground truth (for references of datasets see Tables S1-3) and the Fowlkes Mallows Index (FMI) was used to compare the clustering methods’ performance with the ground truth. For clusterdp the dc parameter was set to 1% and 2% (legend in x axis). (*A*) The number of clusters were set automatically by auto clusterdp (see Fig. S1 for details) and in the case of clusterdv by using the simplex, onion or SI jump criteria. (*B*) The number of clusters were set by choosing the cut-off that corresponds to the known number of clusters, after sorting the cluster centers by the highest γ (clusterdp) or SI values (clusterdv). Grey points in plots are FMI values of each data set. Boxplot indicates median with 25th and 75th percentile hinges, and whiskers extending to the smallest/largest value no less/more than 1.5 × interquartile range from the median. Orange line marks the median. Paired Mann–Whitney test corrected for multiple comparisons by Holm-Bonferroni method, n = 31, comparisons with p < 0.05 were labeled in plots.

### Clusterdv is able to categorize zebrafish startle behavior

The aim of clustering analysis is to identify groupings on real-world data that correspond to real natural phenomena. We used clusterdv to perform unsupervised categorization of movements of zebrafish larvae responding to acoustic startles. Larvae, which swim in short bouts of movement, execute two types of escapes (C-starts) in response to acoustic startles: one bout type with short latency (SLC) and another with long latency (LLC) (Fig. 5*A*) [14]. Critically, both responses differ in a set of kinematic parameters and the neural circuits that produce these behaviors (Fig. 5*B*, left panel and Fig. S9) [14]. The proportion of both C-starts that fish execute varies with the experimental conditions, creating data sets with uneven distributions of bouts. Also, the fish do not always respond to the stimulus with C-starts, so these data sets often have bouts that appear in kinematic space as outliers or that degrade cluster boundaries. In spite of these challenges, the SLC responses are stereotypical and form a tight cluster that appears distinct from the wide spread cluster that corresponds to the LLC bout type (Fig. 5*B,* left panel). These two swim types are relatively easy to categorize by sorting them by the response latency to stimulus onset (red line in Fig. 5*A*) or by setting a threshold that separates them in kinematic space (green line in left panel of Fig. 5*B*). If swim bouts of each category are superimposed it becomes apparent that they correspond to two types of movements that are stereotypical within group (Fig. 5*B*, middle and right panels).

**Fig. 5.**
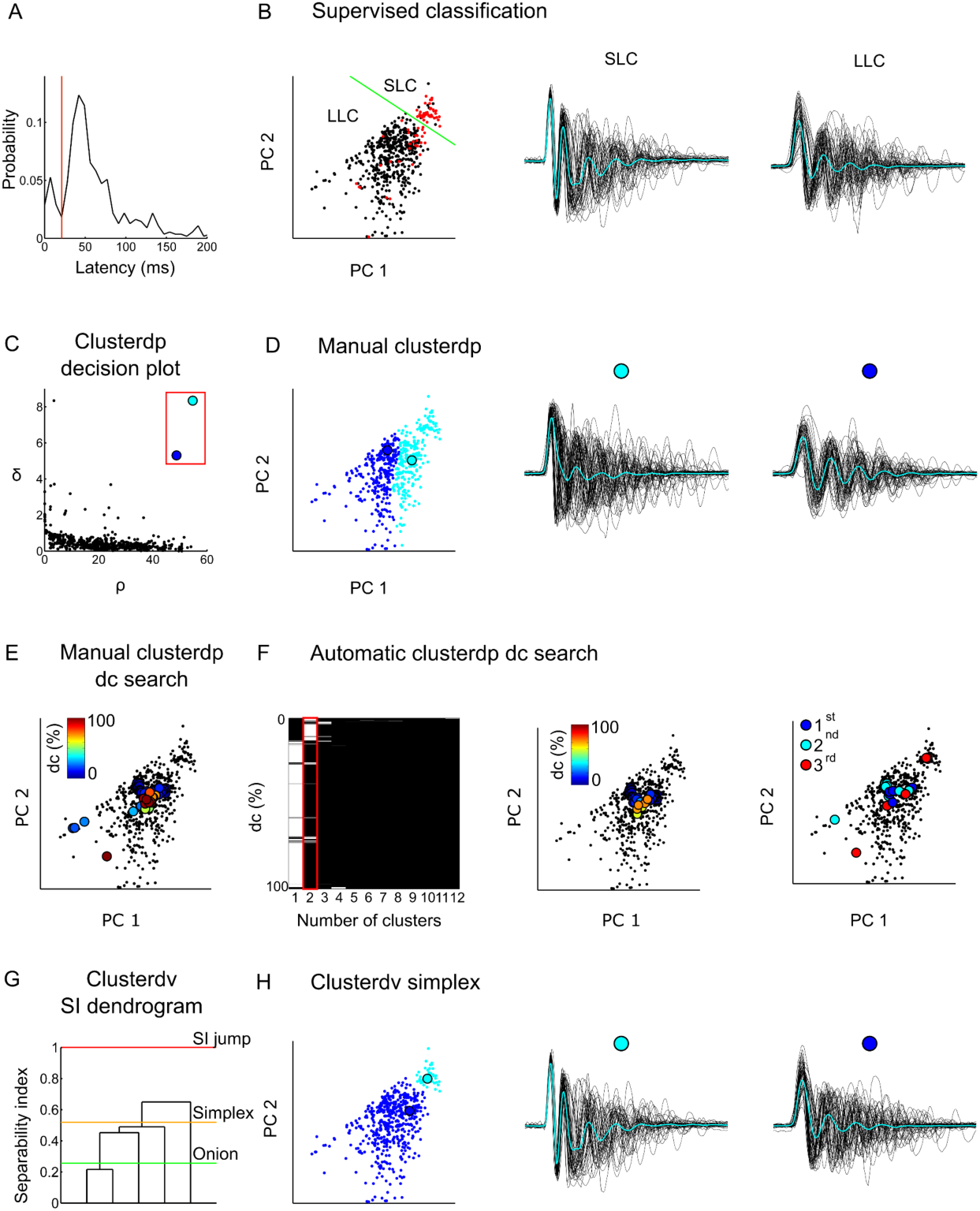
Clusterdv enables unsupervised categorization of zebrafish acoustic startle behavior. (*A*) Swim bout latency from start of acoustic startle. Red line divides swims bouts with short latency (SLC) from bouts with long latency (LLC). (*B*, left panel) PCA applied to swim bout kinematic parameters. Red points correspond to short latency bouts, while black points correspond to long latency bouts categorized by red line in (*A*). Green line separates the small SLC cluster from the larger LLC cluster. Angle of caudal tail segment (°) versus time (ms) of fifty randomly picked SLC (middle panel) and LLC bouts (right panel) categorized by manually dividing the kinematic space (right panel). Cyan lines are the average of all bouts in each category. (*C*) Clusterdp decision plot (ρ vs δ) of bout distribution in (*B*). (D, left panel) Clusterdp assignment of bout distribution in (*B*) after manually picking cluster centers (red rectangle in (*C*)). Angle of caudal tail segment (°) versus time (ms) of fifty randomly picked bouts from solution obtained in (*C*). (*E*) Locations of cluster centers in data picked manually by humans for different dc values. Colors represent dc values according to legend. (*F*) Clustering solutions for automatic clusterdp as a function of dc values (left panel). Red outline shows the known 2-cluster solution. Locations of cluster centers in data set obtained from automatic clusterdp solutions for two cluster solutions (*middle panel*) and the three highest ranking cluster solutions (right panel). Colors in middle panel represent dc values as in legend and in right panel indicate the γ ranking. (G) Clusterdv SI dendrogram for data in (*B*). Lines represent cut-off criteria used to choose number of clusters. (*H*, left panel) Clusterdv assignment according to the simplex threshold in (*G*). Angle of caudal tail segment (°) versus time (ms) of fifty randomly picked SLC (middle panel) and LLC bouts (right panel) categorized according to the simplex cut-off (orange line in (*G*)).

We applied clusterdp to this data set and found that it consistently failed to find the 2 correct clusters (Fig. 5*C-D*). As with the “exclamation mark 1” data set the clusterdp rule fails to rank the cluster centers correctly, not identifying the small SLC cluster center before splitting the larger LLC into several clusters, regardless of the choice of dc or the criteria to select the cluster number (Fig. 5*E-F*). When we applied clusterdv to this data set, the cluster center ranking was correct (Fig. 5G), and the algorithm was able, using an automatic criterion, to find essentially the same solution that was found by drawing a line in kinematic space or sorting the swims by latency (compare Fig. 5*B* with Fig. 5*H*). Clusterdv is thus able to categorize automatically animal behavior in a data set with uneven sparse data that is corrupted by noise.

## Discussion

We have described here a novel, robust and simple density-based clustering algorithm, clusterdv, based on the density valleys between data points, that is applicable to a wide variety of data. It delivers better results than clusterdp, the state of the art density-based clustering algorithm [6]. In particular, it is able to find tight clusters of low density, where clusterdp fails, because its rule gives more importance to gaps between clusters than distances. Thus, a fully automatic version of clusterdv significantly outperforms clusterdp across a wide variety of data sets, even in the case where the number of cluster centers is set for clusterdp based on prior knowledge of the data structure (Fig S10).

All density-based clustering methods suffer from the problem that density estimation for data with finite sample size produces “sporadic” local maxima that are not related to the “real” structure present in data. To deal with this problem clusterdv produces a hierarchical tree of “putative” cluster centers and uses an intuitive metric, the separability index or SI to rank their importance. The number of “putative” cluster centers is often small, because all data points with negative SI values, which are not separated by density valleys, are *a priori* excluded from being cluster centers. To determine how many clusters exist in the data, it is necessary to decide which cluster centers are likely to be genuine, and which may occur sporadically due to sampling error, by setting an appropriate threshold on the SI value, or to cut the SI dendrogram at a particular set of nodes. We automated this step of the algorithm by developing data-based criteria to choose the number of clusters. One such criteria, selection of the largest jump in SI, correctly determined the number of clusters in 29 out of 33 data sets, and is close to the correct solution in all cases, giving results that are comparable to setting the correct number of clusters by hand. It should be noted that this method is based on the assumption that any clustered organization is clearly distinct from noise in the data. For real world data sets, it is not clear that this assumption will always be met. Therefore, to be applied in real-world data sets, such as the zebrafish swimming data we have described here, we developed two other criteria, termed “onion” and “simplex”, similar to the gap statistic method [15], that construct, from the original data, reference distributions that only have one density peak. These reference distributions are used to measure the probability that “sporadic” cluster centers may arise in that particular set of data. Often the “onion” and “simplex” methods also gave the correct solution for many of the data sets we tested (14 and 21 correct data sets respectively), but other times these methods overestimated the number of clusters. Even when the number of clusters did not match the ground truth, these methods still showed good performance, because the dendrogram reflected the true underlying data structure. It is likely that some of the data sets, in particular the real-world ones, contain other clustered organization that is not captured by the manual labeling. Thus, it is possible, in some cases, that the reference-based methods are uncovering meaningful structure.

Most clustering methods need one, or several, parameters to be set, so that correct results are obtained for different data distributions. These criteria introduce a subjective step in clustering analysis that may impact the particular solutions obtained. This is not the case for clusterdv. We benchmarked clusterdv on a set of 33 distributions with known ground truth that were chosen because they offer difficult challenges to clustering analysis such as: arbitrary cluster shape [16-19], number [20] and spatial distribution [21], clusters with fuzzy edges [22,23], data with multiple dimensions [24,25], corruption with noise [6,26], and distributions with uneven proportions of clusters; and did not need to adjust any parameter to solve a particular data set. Nevertheless, there are parameters that can be set in clusterdv, if desired. One such parameter, the number of divisions used to calculate the density lines, allows a trade-off of computational time vs accuracy, and needs to be set sufficiently large not to degrade the results. The density estimation by Gaussian mixture [8] may be performed using distinct methods or rules, but we found that the one simple heuristic used here always gave satisfactory results. Other methods have been proposed that improve on, or automate, aspects of clusterdp, (e.g. [7,27,28]), but, to our knowledge, none has been demonstrated to work well across a similar range of data sets without parameter tuning.

Clusterdv does not work directly with distance-based data, because it is necessary to embed this kind of data in a low dimensional space to calculate ρ and the density paths. Since our aim was to benchmark clusterdv across many types of data and compare it with clusterdp, we choose a commonly used method to reduce dimensionality, t-SNE, that created good low dimensional embeddings for the data sets that we tested, as well as many other types of data [29-31]. Nevertheless, any other method to reduce dimensionality could be combined with clusterdv. As an example, we also applied UMAP [32] to the 4 high dimensional data sets that we used in this work and, although UMAP did not produce, for these particular cases, better feature spaces than t-SNE, clusterdv, in most cases, still outperformed clusterdp with this method (Table S4).

Finally, we applied clusterdv to the difficult problem of unsupervised behavioral categorization. We created a zebrafish larvae behavioral data set that is sparse and composed of highly uneven clusters that are plagued with noise, but is known to contain two distinct swim categories [14]. Clusterdv could identify, in a completely automatic fashion, meaningful behavioral categories that these animals use when startled with acoustic stimuli, while clusterdp failed to provide correct results.

In many situations, it is important to determine the clusters that exist in a data set, without a priori knowledge of their number or shape. To do this with confidence requires a method that delivers consistent results, and robustly selects the correct number and distribution of clusters. The systematic validation of clusterdv across many artificial and real world data sets, makes it suitable to apply to novel problems. We expect clusterdv to be useful in analyzing a wide range of data that has structure that reflects natural phenomena, but where the ground truth is unknown.

## Materials and Methods

### Animal care

Fish were reared on a 14/10 hour light/dark cycle at 28 °C on sets of 20 in E3 water as described previously [33]. Wildtype Tübingen zebrafish larvae (6-7 days post fertilization) were used for behavioral experiments. Animal handling and experimental procedures were approved by the Champalimaud Foundation Ethics Committee and the Portuguese Direcção Geral Veterinária and were performed according to the European Directive 2010/63/EU.

### Behavioral assays

Behavioral recordings were performed in acrylic transparent 5 × 5 cm arenas in groups of 7 fish. 100 ms. tones with fixed frequency (200 to 2000 Hz) where delivered, every 2 min, by two HP mini speakers (Hewlett-Packard Company) attached to the arena. The fish’s movements were recorded at 700 frames per second using a custom made behavior tracking system composed of a high-speed infrared sensitive camera (MC1362, Mikrotron) fitted with a Schneider apo-Xenoplan 2.0/24 lens. Fish were illuminated by a 10 × 10 cm LED-based diffusive backlight (850 nm, Nerlite) placed below the fish arena and the camera was fitted with a 790 nm long pass filter that blocked visible light. Fish and tail tracking were performed online by custom written programs using C# (Microsoft) and the OpenCV imaging processing library as described previously [34]. Kinematic parameters and swim bout detection were performed using custom written Matlab made programs [35]. A subset of 550 bouts was randomly picked to construct the “beep20” data set.

#### Density valley clustering algorithm

The local densities are estimated from a KDE (Kernel density Estimate) calculated using Gaussian kernels, using a Matlab-based toolbox developed by Alexander Ihler (http://www.ics.uci.edu/∼ihler/code/kde.html). The bandwidth for each data point was chosen using a leave-one-out maximum likelihood estimate, with all the bandwidths constrained to be proportional to the kth nearest neighbor distance where:

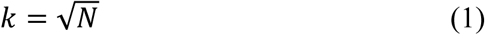

and N is the number of data points. We calculate the density valley depth between points by using the KDE to estimate the density at fixed intervals along straight lines connecting pairs of points and finding the minimum value along each line. We limited the computational time of this step for larger data sets, by restricting the calculation to three sets of edges: 1) the minimal spanning tree found using Kruskal’s algorithm [36], 2) edges connecting each point to the nearest point of higher density, and 3) edges connecting each point to the nearest 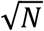points. To identify the smallest drop in density on any path connecting two points, we applied the single link algorithm to the matrix of values [9]. Based on that value we calculate the separability index (SI) for each point by:

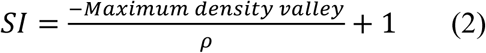

where ρ is the local density at that point. Points with positive SI values represent putative cluster centers, with values ranging from 0 (not separated from other cluster centers) to 1 (completely separated from other clusters).

#### Generation of reference distributions

We used two methods to generate reference distributions that aimed to match features of the original distributions, but should only have a single density peak. The “simplex” method aims at creating a reference distribution within the same convex hull as the original data but without multiple density peaks. Briefly, every set of D + 1 points in the data set, where D is the number of dimensions of the data, defines a simplex. We sample each reference point randomly from within one of these simplexes, which is itself chosen randomly with a weighting proportional to its volume. This has the effect of sampling each simplex at equal density, resulting in a summed distribution which has only one peak. This produces a distribution that lies within the convex hull of the original data, and therefore has a similar range of pairwise point distances. However, the range of local point densities can be very different, which may not be ideal for the clusterdv method, for which these densities define the x-axis range of the decision plot. Therefore, for the “onion” method we aim to match not the shape, but the distribution of point densities in the original data. We make the “onion” reference distribution by first calculating the local density at each of the original data points, and arranging them in descending order. We then choose the first point of the reference distribution randomly from within an n-sphere, whose radius is chosen so that the local density matches that of the densest point. Subsequent points are then chosen from n-spherical shells surrounding the previous level, whose outer radii are chosen to match the local density within each shell with the density of the corresponding point in the original distribution. The “onion” method creates a spherically symmetrical reference distribution with a single local maximum, and a distribution of density that is very similar to the original data.

#### Determination of number of clusters for clusterdv

Data points with positive SI value were considered ‘putative’ cluster centers. Their SI value was used to construct dendrograms that reflect the ranking of cluster centers. As cluster centers with decreasing SI are added to the dendrogram, they are connected to the cluster center with which they co-partition at the next higher level. For all data sets we computed three different criteria for determining a threshold SI value at which to cut the dendrogram. For the ‘max SI jump’ we take the largest jump in the separability index (SI jump) between successive cluster centers. Alternatively, we take the 95th percentile of the second highest SI value from 100 reference distributions computed using the “simplex” or the “onion” methods. All cluster centers with higher SI values than each of these criteria formed the associated solutions for each data set.

#### Automatic cluster center identification for clusterdp

The local densities (ρ) and δ value where calculated for every data point as described in [6]. By definition γ_i_ = ρ_i_δ_i_ will be large for cluster centers that belong to clusters that are well separated and have many points. We rely on this quantity to find a “jump” on the data that signals when data points stop being cluster centers and start being data points belonging to an already identified cluster, or outliers. 100 unimodal reference distributions using the “simplex” resampling method were created. This method was used to create unimodal distributions because clusterdp uses pairwise distances to calculate its two measures and the distances between points are highly influenced by the shape of the distribution. Next, we calculate the γ measure for the real data and the reference distributions, sort these values, average the γ sorted values of the reference distributions and subtract those from the γ sorted values of the original data. To find the appropriate number of clusters we determined how much the γ difference values deviate from the reference distributions. To determine this, the γ difference values are divided by the values of the original distribution with all values made positive by subtracting the minimum (normalized γ differences). The cluster centers were found by choosing the points before the largest jump in the normalized γ values.

#### Analysis of the MNIST data set

The test data of the MNIST digit data set was placed in a set into a three-dimensional space obtained by parametric t-SNE (perplexity = 30) [37]. For manual selection of the SI cut-off, two methods were used. First the tree was cut at a level which gave 10 clusters. Alternatively, the ‘balanced’ cut was found by starting with the 10 cluster solution and lowering the threshold on the branch with the largest number of points, until another center was found, and pruning the branch with the smallest number of points. This process was repeated until the standard deviation of cluster sizes was at a local minimum. The resulting cluster assignments were then used to classify the 60000 remaining digits via a nearest neighbor method.

#### Dimension reduction of data

Both the UMAP [32] and t-SNE [38] methods were used to reduce the dimensionality of: bone marrow [22], Zachary [25], Olivetti [11] and MNIST test [12] data sets. The specific parameters for each data set are displayed for the t-SNE in Table S1 and for UMAP in Table S4.

#### Data point assignment

For all data sets the non-cluster center points were assigned, sequentially and in decreasing density order, to the same cluster of the nearest neighbor of higher density [6].

#### Automatic clusterdp validation

To determine the effectiveness of the automatic clusterdp we compared the algorithm’s success in finding the correct number of clusters to the ability of people manually picking the cluster centers. d_c_ values from 1 to 100 % were applied to nine synthetic data sets and their decision plots (ρ vs δ) were shown to 9 unpaid volunteers by using a custom made Matlab script (MathWorks). The participants were asked to pick the cluster centers that they thought were present in the decision plots by drawing a square around them. The order of display of the decision plots was randomized. The people were not given feedback over their choices or informed of the number of clusters present in the data sets. To each person it was explained how the clusterdv algorithm works and how to draw a square using the Matlab custom program (MathWorks). For comparison of two independent groups, significance of difference was tested with the Mann–Whitney test. Statistical analysis was performed using the Statistic Toolbox (MathWorks). Differences were considered significant when *P* < 0.05.

## Figure Legends

**Fig. S1.**
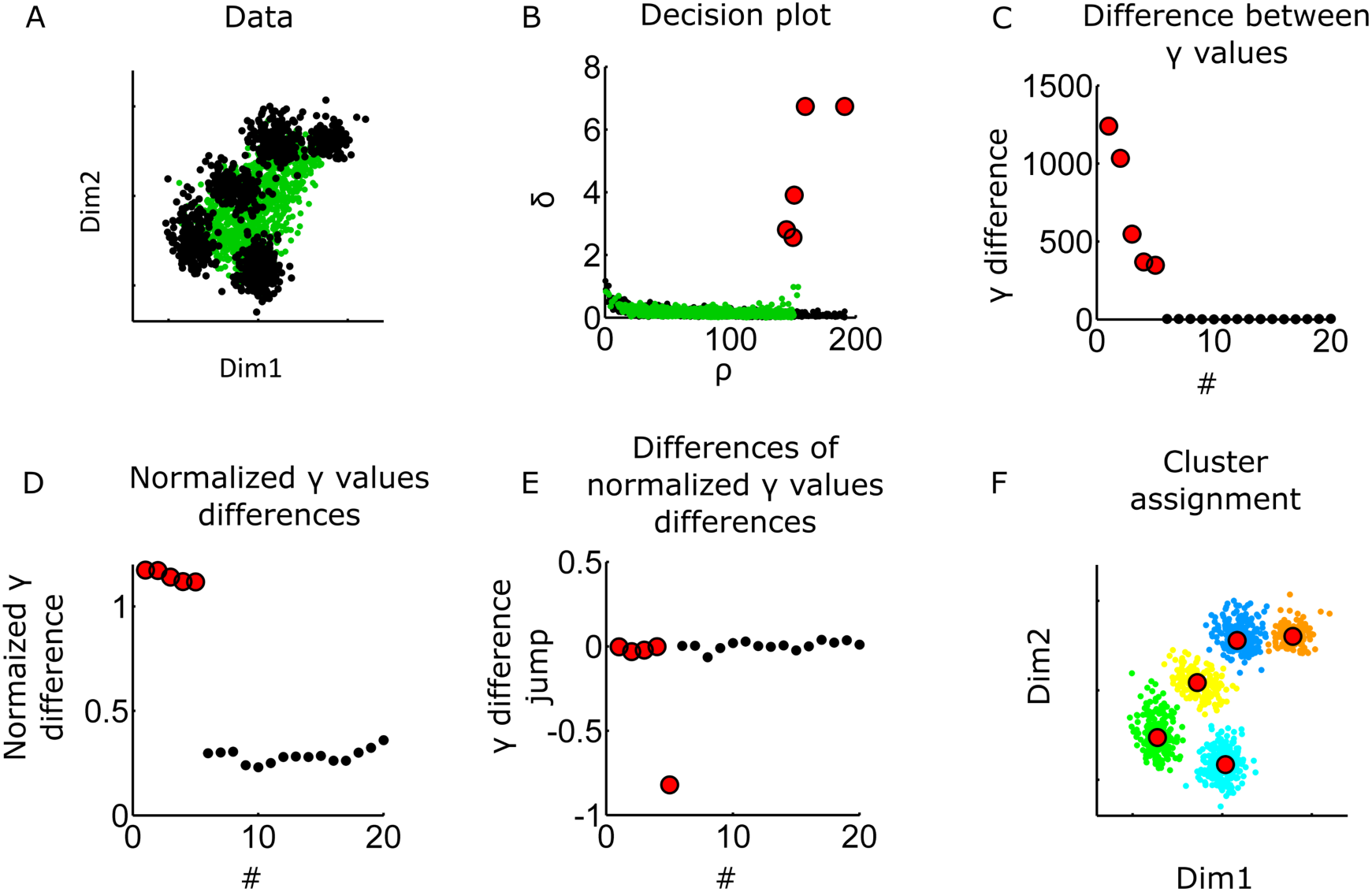
Automatic clusterdp method. Automatic method to determine the number of cluster centers for the density peak algorithm (see Materials Methods for details) [6]. (*A*) Point distribution drawn from a mixture of five gaussians (black). In green, an example single peak reference distribution obtained by resampling the distribution in (*A*) using the “simplex” method. (*B*) Decision plot (δ vs ρ) of the two distributions in (*A*). Black and green points correspond to distributions in (*A*). (*C*) Difference between the γ (δ*ρ) values of the gaussian mixture distribution and the average values of 100 reference distributions, the points are ordered for decreasing values of γ difference values. (*D*) Normalized γ difference values. (*E*) Difference of values in (*D*), or “jump”. The minimum point is used to choose the number of cluster centers. (*F*) Point assignment of black data set in (*A*) according to the number of clusters found in (*E*). The five clusters found in (*F*) correspond to the five gaussians that form the (*A*) distribution. Red points correspond to cluster centers found by the algorithm.

**Fig. S2.**
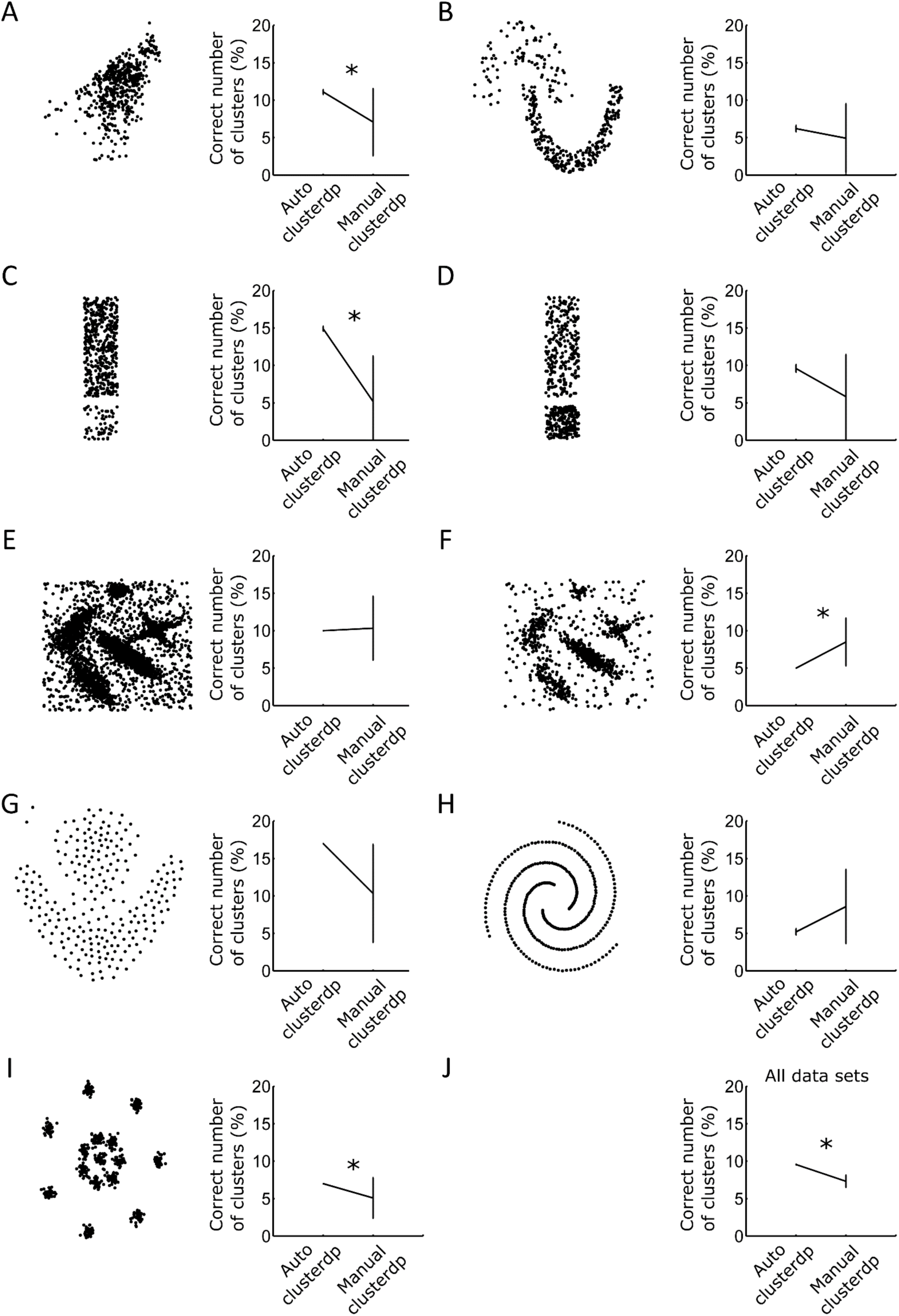
Automatic clusterdp performance. We compared the performance of automatic clusterdp to pick the correct number of clusters to the human ability to manually pick cluster centers (see Materials and Methods for details). (*A, C*-*D*) This study, (*B*) [18], (*E*-*F*) [6], (*G*) [17], (*H*) [19], (*I*) [21], and (*J*) all data sets together. Error bars are standard error of the mean. Mann–Whitney test, * p < 0.05, auto n = 10, manual n = 9.

**Fig. S3.**
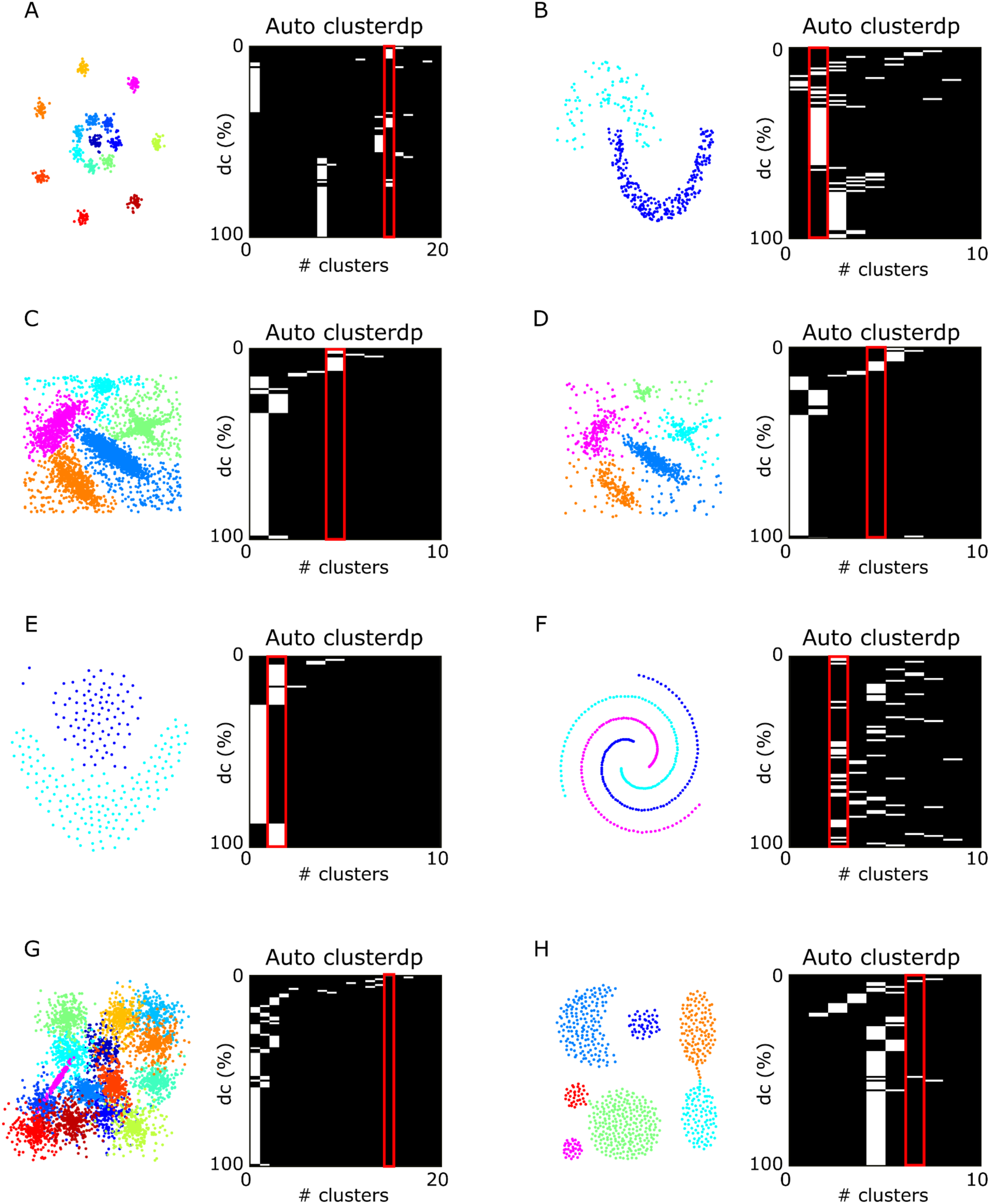
Accuracy of the clusterdp algorithm is critically dependent on the choice of parameter dc. Left panels represent synthetic point distributions used in [6]: (*A*) [21], (*B*) [18], (*C*-*D*) [6], (*E*) [17], (*F*) [19], (*G*) [23], and (*H*) [16]. White lines in right panels correspond to number of clusters found using automatic clusterdp for a particular d_c_ value expressed in percentile of all pairwise distances. The red outlines mark the column with the correct number of clusters.

**Fig. S4.**
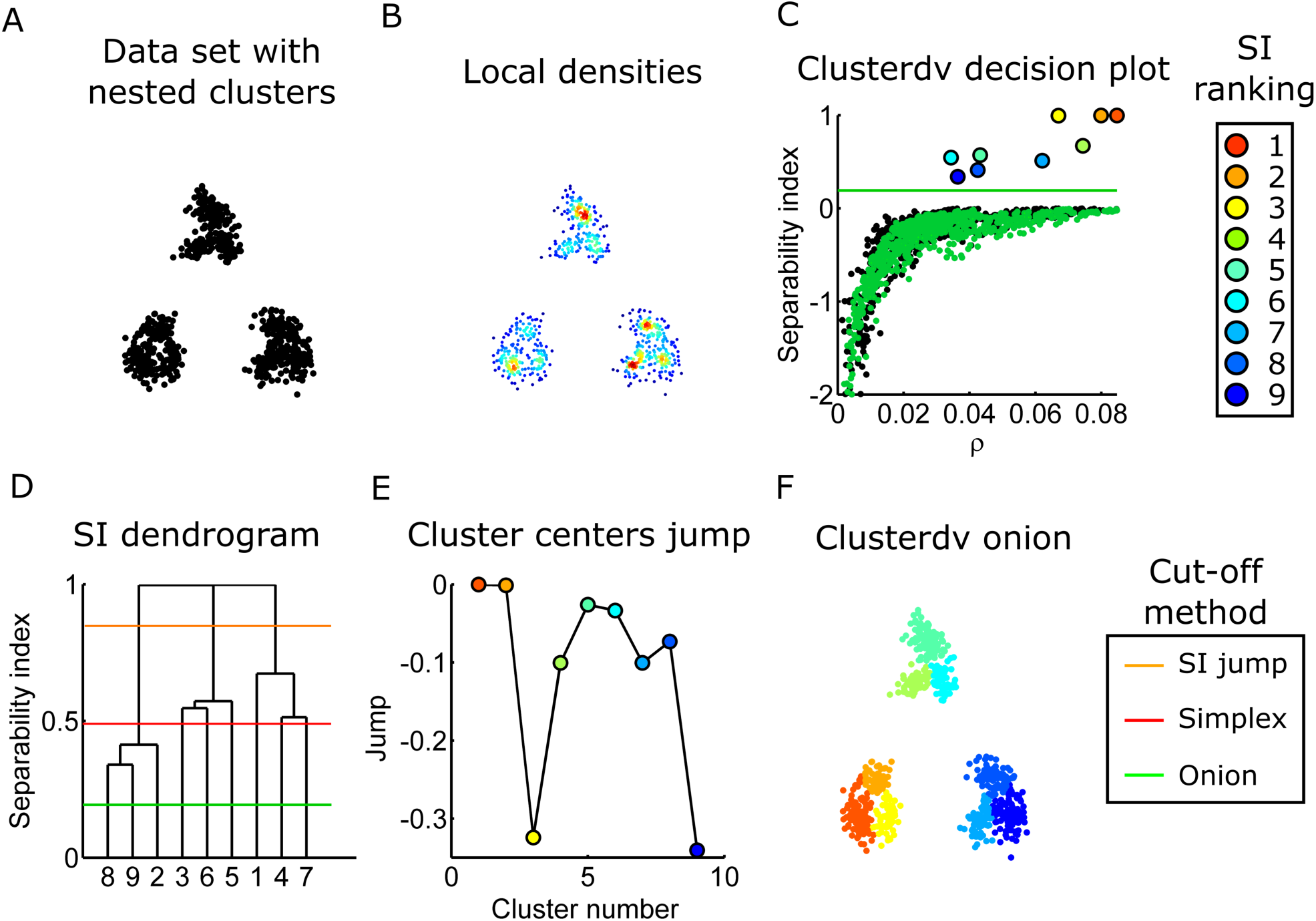
Solution of clusterdv for data set with nested groups of clusters. (*A*) Point distribution drawn from mixture of nine gaussians. (*B*) Local densities are calculated using adaptive gaussian density estimator. (*C*) SI versus local density (ρ). Green points are from example reference distribution calculate using the onion method. Circles represent cluster centers and their colors represent their ranking according to the SI value (see legend on the top right). (*D*) SI dendrogram of (*C*). Lines represent the cut-off method used to choose number of clusters (see legend in the bottom right). (*E*) SI jump versus cluster centers sorted by decreasing SI value. Colors of circles as in (*C*). (*F*) Cluster assignment of the point distribution in (*A*) using the onion cut-off.

**Fig. S5.**
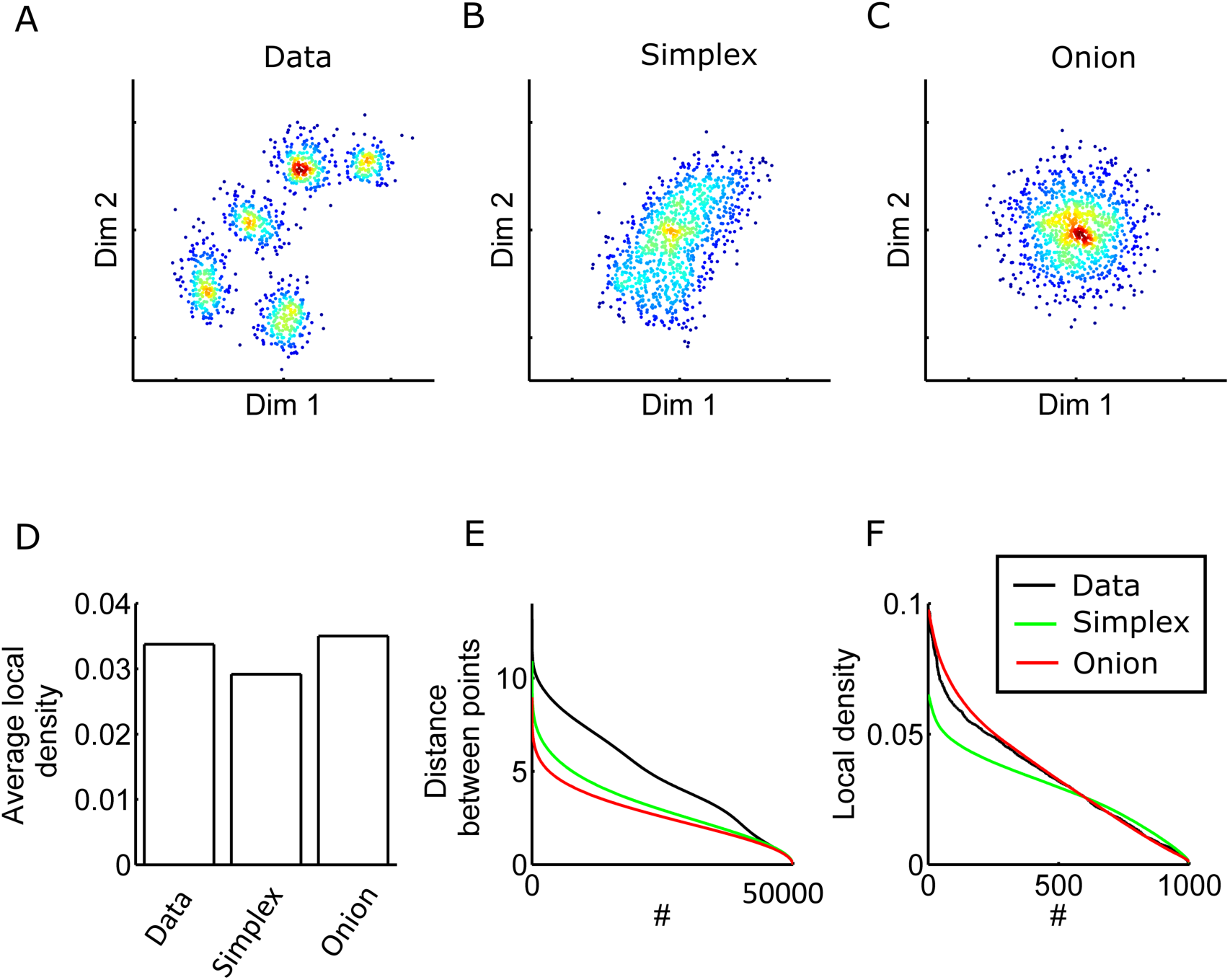
Methods to calculate reference distributions. (*A*) Synthetic point distribution drawn from a mixture of five gaussians. (*B*) Reference distribution obtained by resampling the distribution in (*A*) using the simplex method. (*C*) Reference distribution obtained by resampling the distribution in (*A*) using the onion method. (*A*-*C*) Colors represent local densities calculated by using an adaptive mixture of gaussians density estimate. (*D*) Average local density between the three distributions in (*A*-*C*). (*E*) Distance between points plotted in descending order. (*F*) Local densities plotted in descending order. (*E*-*F*) Colors of lines represent distribution types according to legend.

**Fig. S6.**
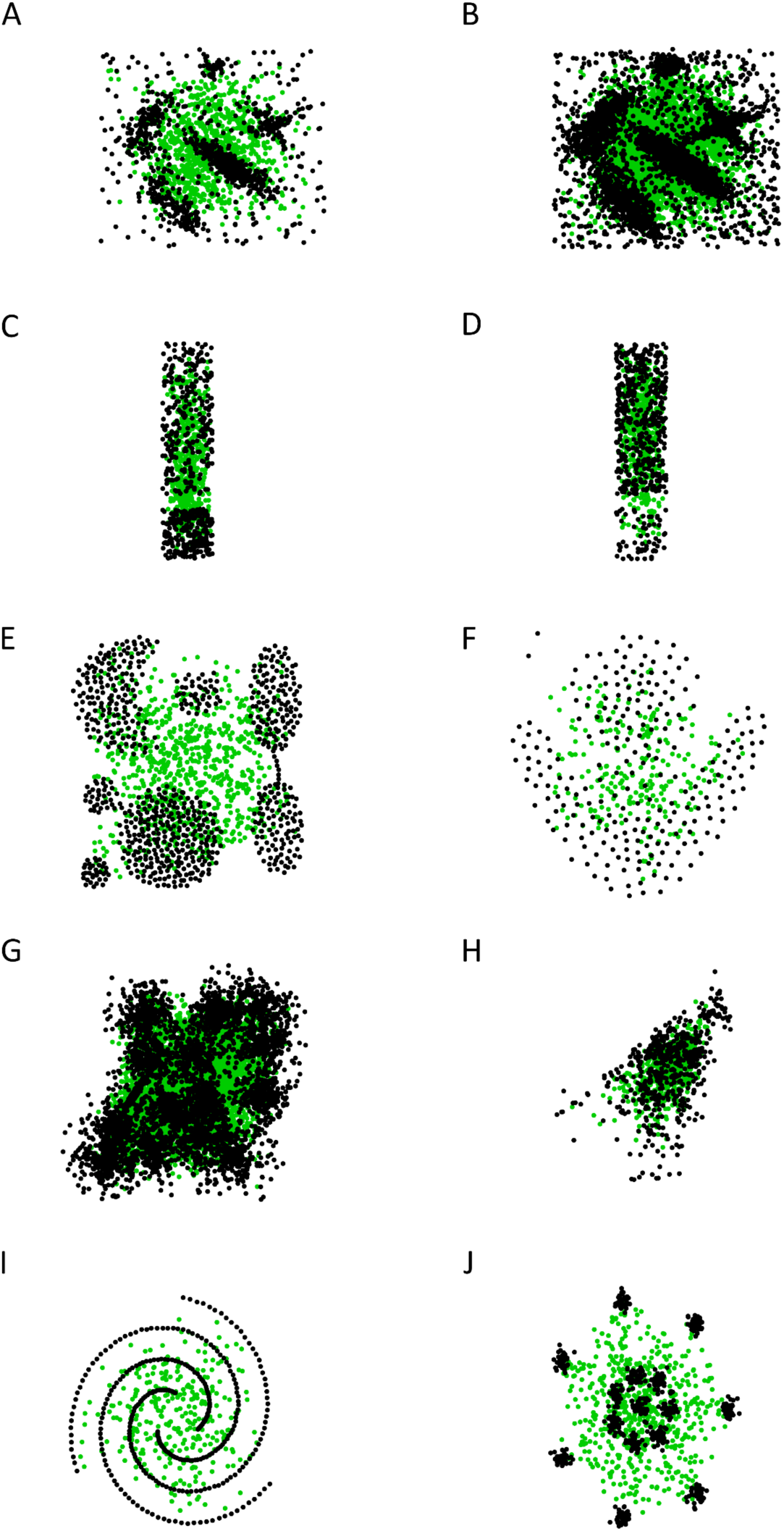
Example reference distributions using the simplex method. Black points are original point distributions. Green points are “simplex” reference distributions. Synthetic point distributions from: (*A*-*B*) [6], (*C*-*D, H*) this study, (*E*) [16], (*F*) [17], (*G*) [23], (*I*) [19], and (*J*) [21].

**Fig. S7.**
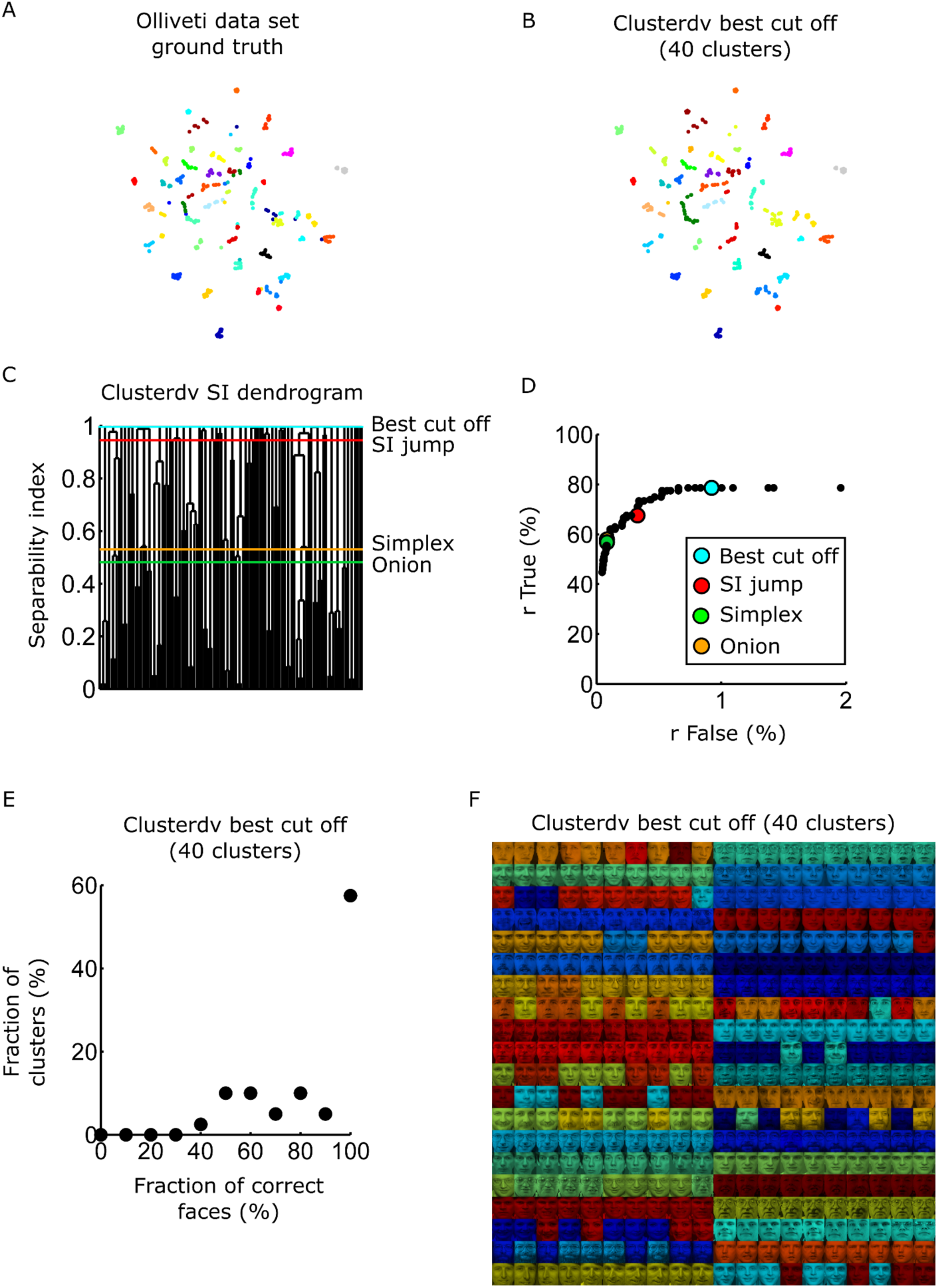
Density valley clustering of the Olivetti face data set. Complex wavelet structural similarity (CW-SSIM) index [11] was applied to the Olivetti face data set [10] to calculate similarities between pairs of images. Since clusterdv is not a distance-based clustering method we applied t-SNE to reduce the dimensionality of the similarity matrix (perplexity 30). (*A*-*B*) CW-SSIM embedded in a two-dimensional t-SNE space. Point colors correspond to ground truth (*A*) and clusterdv solution of the best cut-off (40 clusters with highest SI value) (*B*). (*C*) SI dendrogram of data in (*A*-*B*). Colors of lines correspond to the cut-off criteria used to choose cluster centers according to legend. (*D*) Fraction of pair of images correctly associated to the same cluster (rTrue) as a function of the fraction of pair of images erroneously assigned to the same cluster (rFalse), using different cut-off values. Colors of circles correspond to the cut-off criteria according to the legend. (*E*) Histogram of percent matching faces for the largest group in each cluster for the solution obtained using the best cut-off criteria (40 clusters). (*F*) Assignment of the Olivetti data set for the best cut-off criteria (40 clusters). Each picture is labeled with the color of the cluster it was assigned to.

**Fig. S8.**
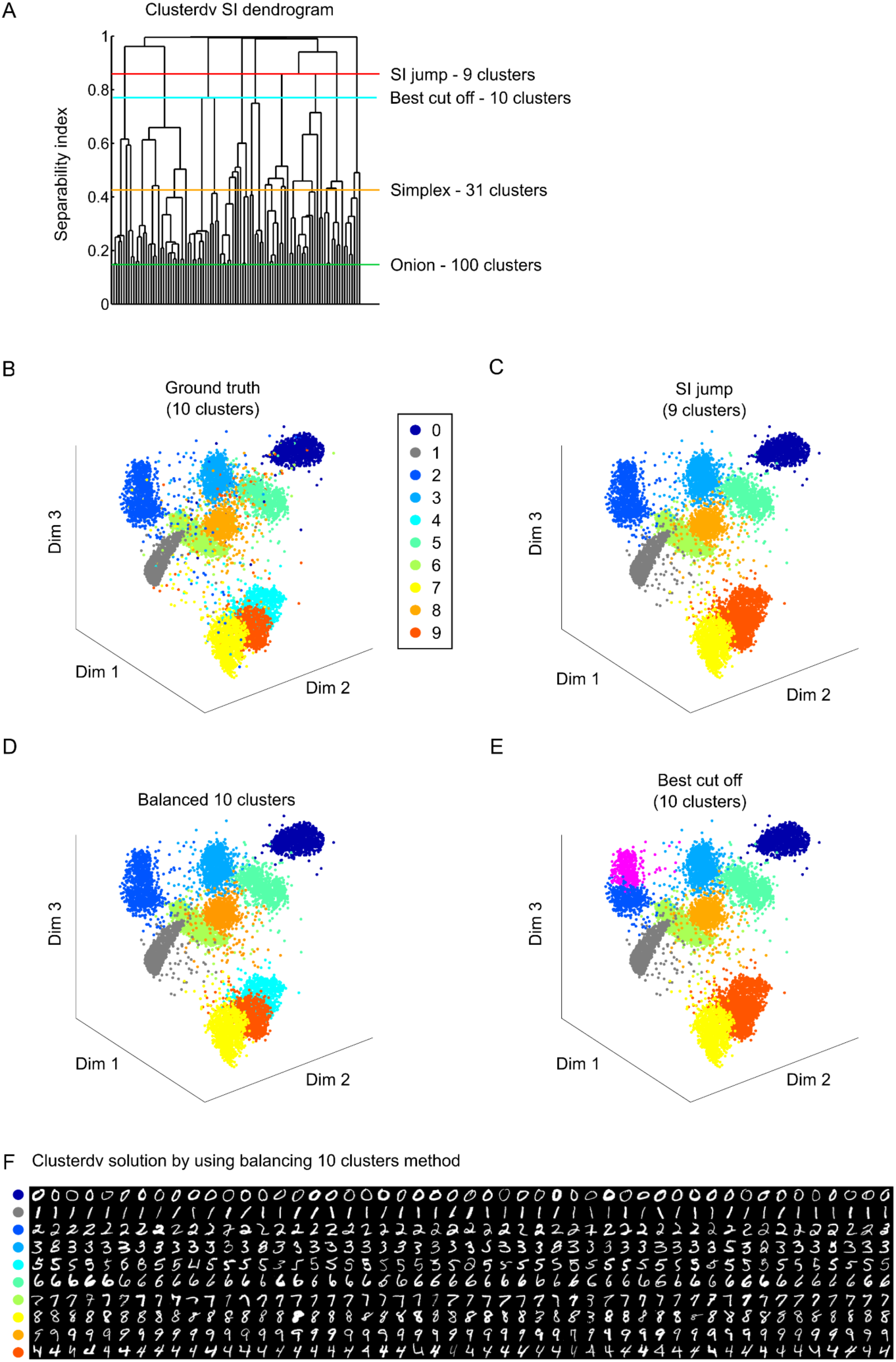
Density valley clustering of the MNIST handwritten digit data set. Parametric t-SNE was used to embed the raw data of the test MNIST data set into a three-dimensional space (perplexity = 30) [37]. (*A*) SI dendrogram of the MNIST test data set. Red line, SI jump. Orange line, simplex threshold. Green line, onion threshold. Cyan line best cut-off (10 clusters with higher SI values). (*B*) Colors represent ground truth according to legend. (*C*) Solution given by SI jump. Most clusters are correct with the exception of clusters of digits 9 and 4 that were grouped together because these clusters are not well separated in the t-SNE space. Compare clusters 4 and 9 in (*B*) with dark orange cluster in (*C*). (*D*) Solution by finding clusters in dendrogram by minimizing standard deviation of number of points per cluster (see Materials and Methods for details). (*E*) Solution given by using best single cut-off (10 clusters). Similar to the SI jump solution but cluster 2 was divided into two clusters (compare cluster 2 in (*B*) with blue and magenta cluster in (*E*)). (*F*) Fifty digits were picked randomly from solution in (*D*).

**Fig. S9.**
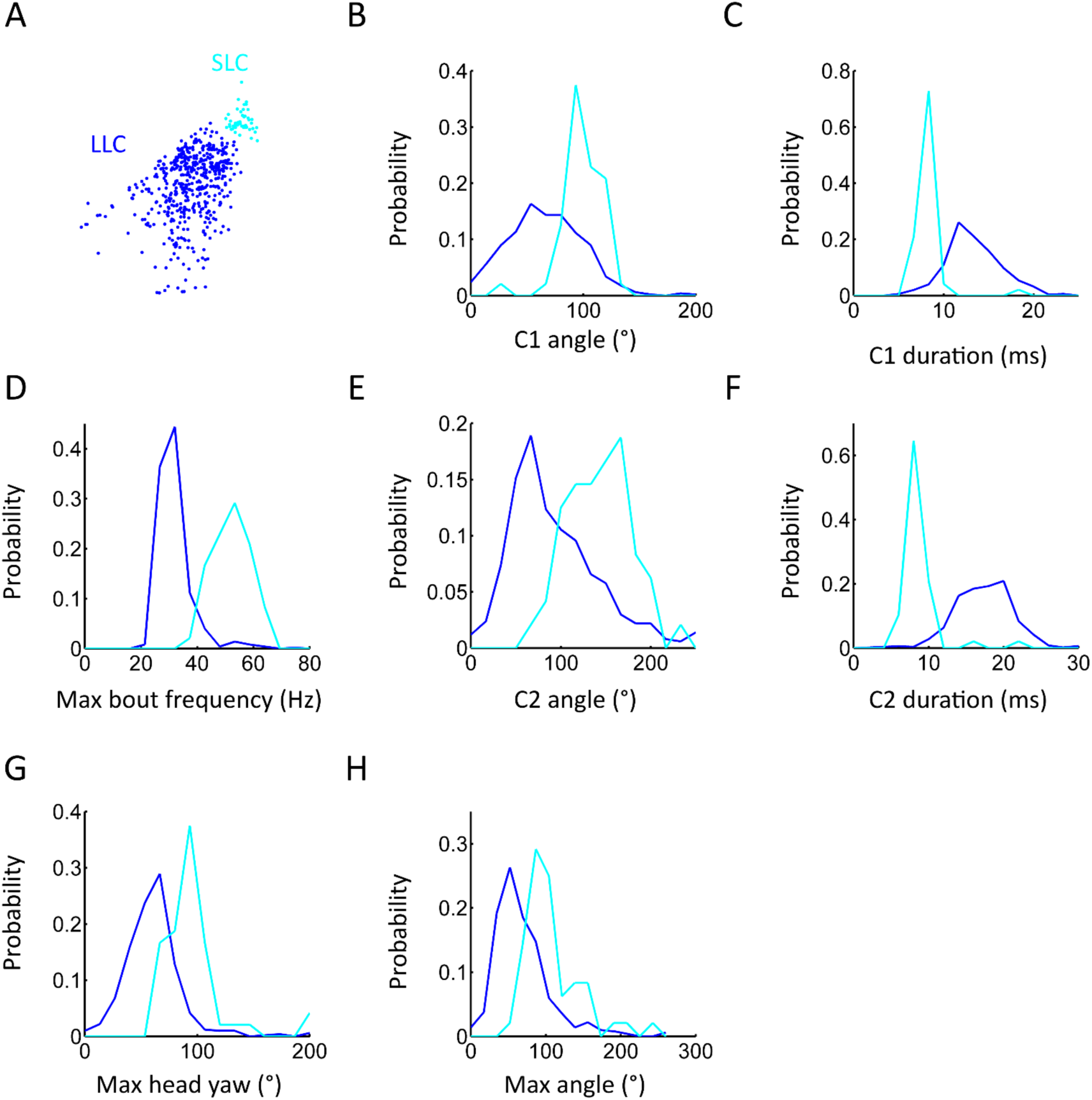
Short latency and long latency bout types have distinct distributions of kinematic parameters. (*A*) Swim bout data set obtained by larvae responding to acoustic stimuli. Colors represent bout categorization obtained by clusterdv (Fig. 5*M*-*P*). (*B*-*H*) Distributions were obtained by using all bouts in (*A*). SLC and LLC bout types show different kinematic parameter values that are in accordance with previous reports [14]. Colors of distributions correspond to clusters in (*A*).

**Fig. S10.**
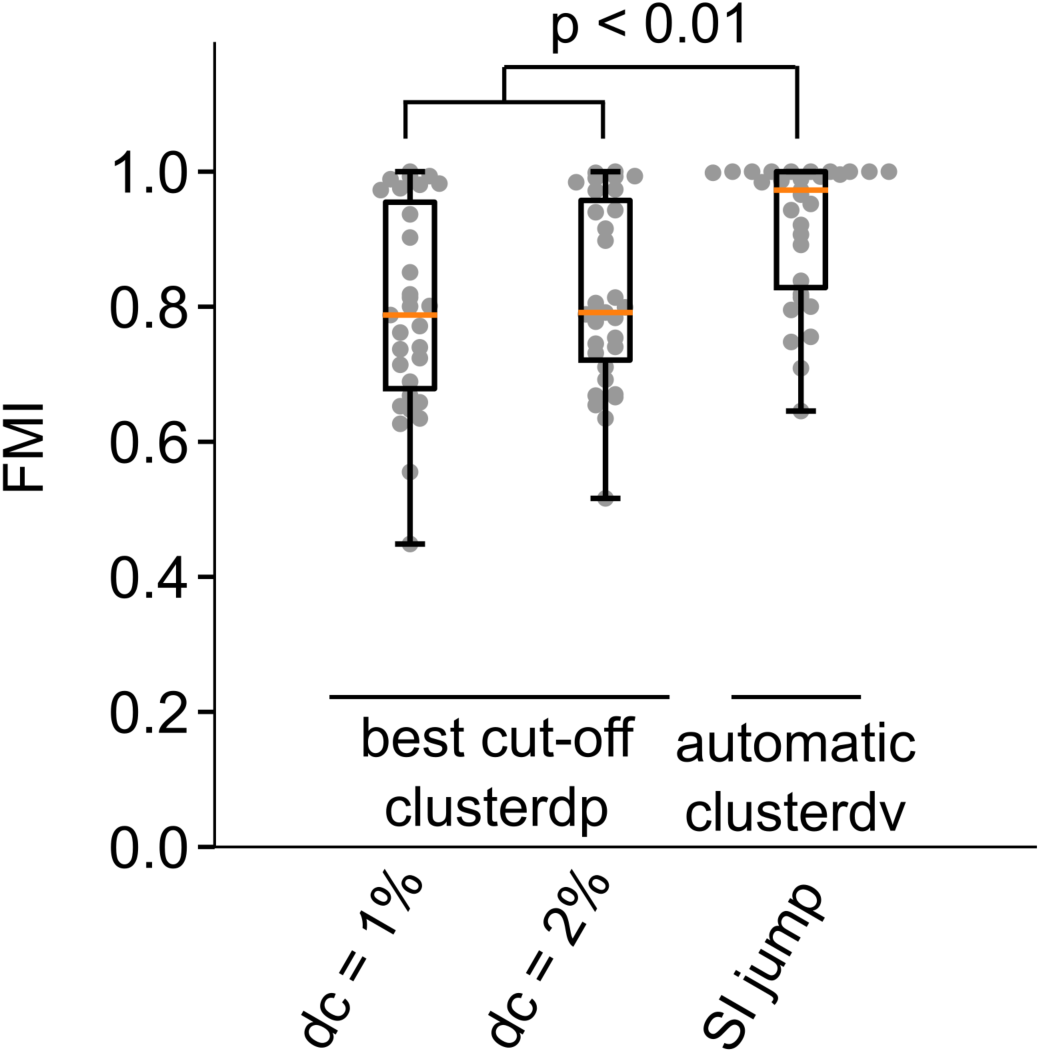
Automatic clusterdv outperforms clusterdp even in the case of cluster centers being set by best cut-off. The clusterdp and clusterdv methods (legends in x axis) were applied to 31 data sets (for references of datasets see Tables S1-3) and the Fowlkes Mallows Index (FMI) was used to compare the clustering methods’ performance with the ground truth. For clusterdp, the dc parameter was set to 1% and 2% (legend in x axis). In the case of clusterdp the number of clusters were set by choosing the cut-off that corresponds to the known number of clusters after sorting the cluster centers by the highest γ (clusterdp). In the case of clusterdv number of clusters were set automatically by using the SI jump criteria. Grey points in plots are FMI values of each data set. Boxplot indicates median with 25^th^ and 75^th^ percentile hinges, and whiskers extending to the smallest/largest value no less/more than 1.5 × interquartile range from the median. Orange line marks the median. Paired Mann– Whitney test corrected for multiple comparisons by Holm-Bonferroni method, n = 31, comparisons with p < 0.01 were labeled in plots. Same data as in Fig.4.

**Table S1.**
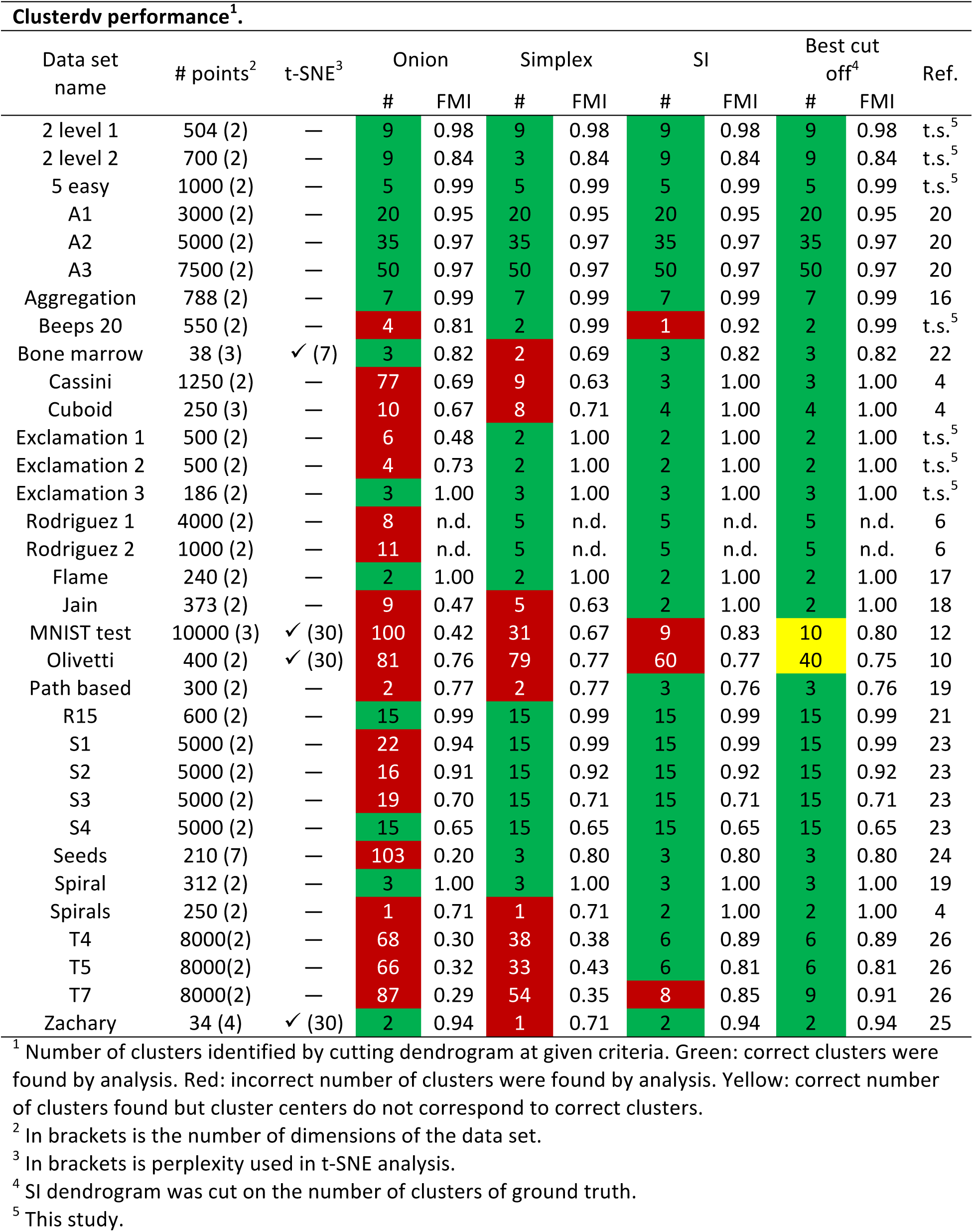
Clusterdv performance.

| Clusterdv performance<sup>1</sup>. |  |  |  |  |  |  |  |  |  |  |  |
| --- | --- | --- | --- | --- | --- | --- | --- | --- | --- | --- | --- |
| Data set name | # points<sup>2</sup> | t-SNE<sup>3</sup> | Onion |  | Simplex |  | SI |  | Best cut off<sup>4</sup> |  | Ref. |
|  |  |  | # | FMI | # | FMI | # | FMI | # | FMI |  |
| 2 level 1 | 504 (2) | — | 9 | 0.98 | 9 | 0.98 | 9 | 0.98 | 9 | 0.98 | t.s.<sup>5</sup> |
| 2 level 2 | 700 (2) | — | 9 | 0.84 | 3 | 0.84 | 9 | 0.84 | 9 | 0.84 | t.s.<sup>5</sup> |
| 5 easy | 1000 (2) | — | 5 | 0.99 | 5 | 0.99 | 5 | 0.99 | 5 | 0.99 | t.s.<sup>5</sup> |
| A1 | 3000 (2) | — | 20 | 0.95 | 20 | 0.95 | 20 | 0.95 | 20 | 0.95 | 20 |
| A2 | 5000 (2) | — | 35 | 0.97 | 35 | 0.97 | 35 | 0.97 | 35 | 0.97 | 20 |
| A3 | 7500 (2) | — | 50 | 0.97 | 50 | 0.97 | 50 | 0.97 | 50 | 0.97 | 20 |
| Aggregation | 788 (2) | — | 7 | 0.99 | 7 | 0.99 | 7 | 0.99 | 7 | 0.99 | 16 |
| Beeps 20 | 550 (2) | — | 4 | 0.81 | 2 | 0.99 | 1 | 0.92 | 2 | 0.99 | t.s.<sup>5</sup> |
| Bone marrow | 38 (3) | ✓ (7) | 3 | 0.82 | 2 | 0.69 | 3 | 0.82 | 3 | 0.82 | 22 |
| Cassini | 1250 (2) | — | 77 | 0.69 | 9 | 0.63 | 3 | 1.00 | 3 | 1.00 | 4 |
| Cuboid | 250 (3) | — | 10 | 0.67 | 8 | 0.71 | 4 | 1.00 | 4 | 1.00 | 4 |
| Exclamation 1 | 500 (2) | — | 6 | 0.48 | 2 | 1.00 | 2 | 1.00 | 2 | 1.00 | t.s.<sup>5</sup> |
| Exclamation 2 | 500 (2) | — | 4 | 0.73 | 2 | 1.00 | 2 | 1.00 | 2 | 1.00 | t.s.<sup>5</sup> |
| Exclamation 3 | 186 (2) | — | 3 | 1.00 | 3 | 1.00 | 3 | 1.00 | 3 | 1.00 | t.s.<sup>5</sup> |
| Rodriguez 1 | 4000 (2) | — | 8 | n.d. | 5 | n.d. | 5 | n.d. | 5 | n.d. | 6 |
| Rodriguez 2 | 1000 (2) | — | 11 | n.d. | 5 | n.d. | 5 | n.d. | 5 | n.d. | 6 |
| Flame | 240 (2) | — | 2 | 1.00 | 2 | 1.00 | 2 | 1.00 | 2 | 1.00 | 17 |
| Jain | 373 (2) | — | 9 | 0.47 | 5 | 0.63 | 2 | 1.00 | 2 | 1.00 | 18 |
| MNIST test | 10000 (3) | ✓ (30) | 100 | 0.42 | 31 | 0.67 | 9 | 0.83 | 10 | 0.80 | 12 |
| Olivetti | 400 (2) | ✓ (30) | 81 | 0.76 | 79 | 0.77 | 60 | 0.77 | 40 | 0.75 | 10 |
| Path based | 300 (2) | — | 2 | 0.77 | 2 | 0.77 | 3 | 0.76 | 3 | 0.76 | 19 |
| R15 | 600 (2) | — | 15 | 0.99 | 15 | 0.99 | 15 | 0.99 | 15 | 0.99 | 21 |
| S1 | 5000 (2) | — | 22 | 0.94 | 15 | 0.99 | 15 | 0.99 | 15 | 0.99 | 23 |
| S2 | 5000 (2) | — | 16 | 0.91 | 15 | 0.92 | 15 | 0.92 | 15 | 0.92 | 23 |
| S3 | 5000 (2) | — | 19 | 0.70 | 15 | 0.71 | 15 | 0.71 | 15 | 0.71 | 23 |
| S4 | 5000 (2) | — | 15 | 0.65 | 15 | 0.65 | 15 | 0.65 | 15 | 0.65 | 23 |
| Seeds | 210 (7) | — | 103 | 0.20 | 3 | 0.80 | 3 | 0.80 | 3 | 0.80 | 24 |
| Spiral | 312 (2) | — | 3 | 1.00 | 3 | 1.00 | 3 | 1.00 | 3 | 1.00 | 19 |
| Spirals | 250 (2) | — | 1 | 0.71 | 1 | 0.71 | 2 | 1.00 | 2 | 1.00 | 4 |
| T4 | 8000(2) | — | 68 | 0.30 | 38 | 0.38 | 6 | 0.89 | 6 | 0.89 | 26 |
| T5 | 8000(2) | — | 66 | 0.32 | 33 | 0.43 | 6 | 0.81 | 6 | 0.81 | 26 |
| T7 | 8000(2) | — | 87 | 0.29 | 54 | 0.35 | 8 | 0.85 | 9 | 0.91 | 26 |
| Zachary | 34 (4) | ✓ (30) | 2 | 0.94 | 1 | 0.71 | 2 | 0.94 | 2 | 0.94 | 25 |

<sup>1</sup> Number of clusters identified by cutting dendrogram at given criteria. Green: correct clusters were found by analysis. Red: incorrect number of clusters were found by analysis. Yellow: correct number of clusters found but cluster centers do not correspond to correct clusters.
<sup>2</sup> In brackets is the number of dimensions of the data set.
<sup>3</sup> In brackets is perplexity used in t-SNE analysis.
<sup>4</sup> SI dendrogram was cut on the number of clusters of ground truth.
<sup>5</sup> This study.

**Table S2.**
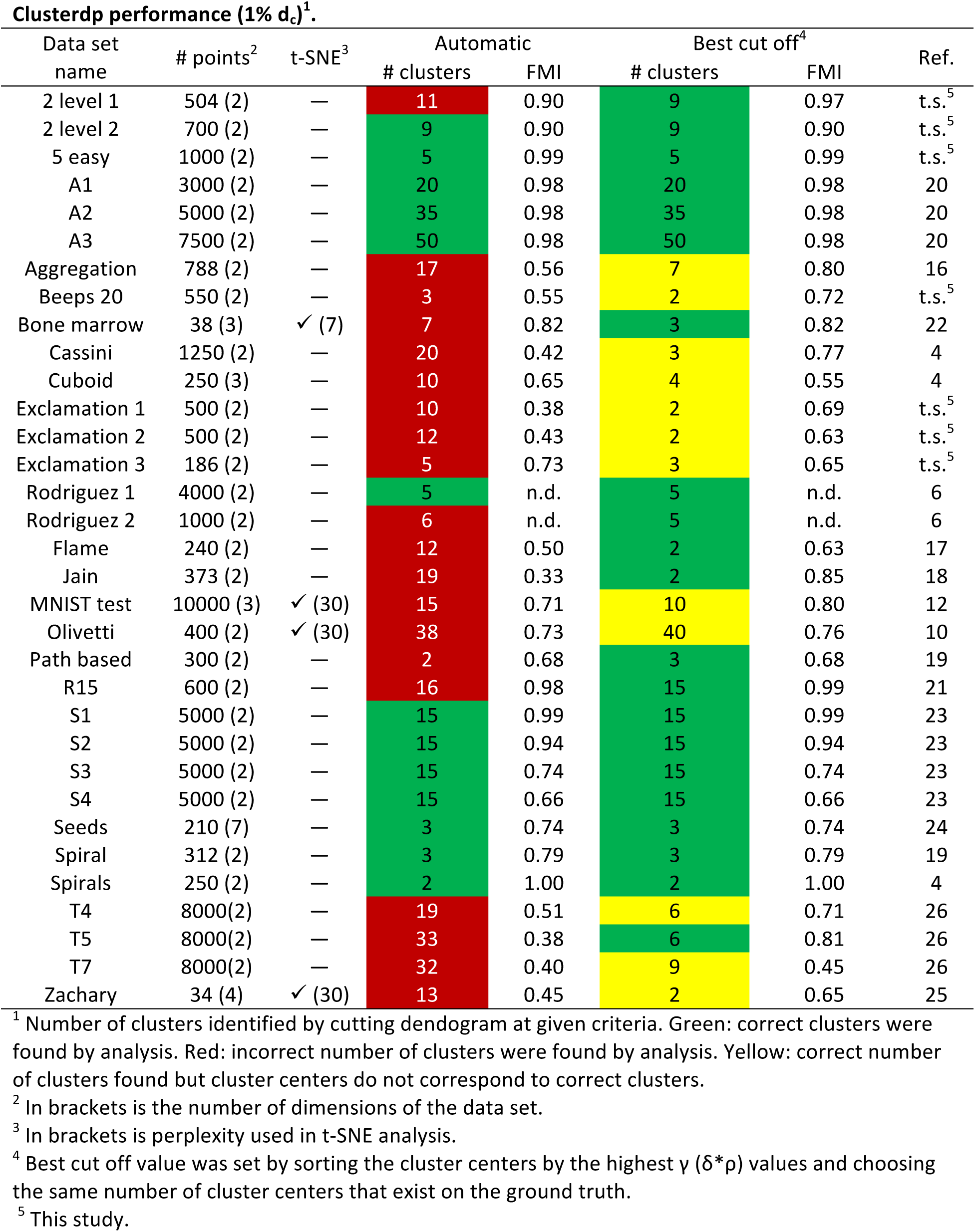
Clusterdp performance at dc 1%.

| Data set name | # points<sup>2</sup> | t-SNE<sup>3</sup> | Automatic |  | Best cut off<sup>4</sup> |  | Ref. |
| --- | --- | --- | --- | --- | --- | --- | --- |
|  |  |  | # clusters | FMI | # clusters | FMI |  |
| 2 level 1 | 504 (2) | — | 11 | 0.90 | 9 | 0.97 | t.s.<sup>5</sup> |
| 2 level 2 | 700 (2) | — | 9 | 0.90 | 9 | 0.90 | t.s.<sup>5</sup> |
| 5 easy | 1000 (2) | — | 5 | 0.99 | 5 | 0.99 | t.s.<sup>5</sup> |
| A1 | 3000 (2) | — | 20 | 0.98 | 20 | 0.98 | 20 |
| A2 | 5000 (2) | — | 35 | 0.98 | 35 | 0.98 | 20 |
| A3 | 7500 (2) | — | 50 | 0.98 | 50 | 0.98 | 20 |
| Aggregation | 788 (2) | — | 17 | 0.56 | 7 | 0.80 | 16 |
| Beeps 20 | 550 (2) | — | 3 | 0.55 | 2 | 0.72 | t.s.<sup>5</sup> |
| Bone marrow | 38 (3) | ✓ (7) | 7 | 0.82 | 3 | 0.82 | 22 |
| Cassini | 1250 (2) | — | 20 | 0.42 | 3 | 0.77 | 4 |
| Cuboid | 250 (3) | — | 10 | 0.65 | 4 | 0.55 | 4 |
| Exclamation 1 | 500 (2) | — | 10 | 0.38 | 2 | 0.69 | t.s.<sup>5</sup> |
| Exclamation 2 | 500 (2) | — | 12 | 0.43 | 2 | 0.63 | t.s.<sup>5</sup> |
| Exclamation 3 | 186 (2) | — | 5 | 0.73 | 3 | 0.65 | t.s.<sup>5</sup> |
| Rodriguez 1 | 4000 (2) | — | 5 | n.d. | 5 | n.d. | 6 |
| Rodriguez 2 | 1000 (2) | — | 6 | n.d. | 5 | n.d. | 6 |
| Flame | 240 (2) | — | 12 | 0.50 | 2 | 0.63 | 17 |
| Jain | 373 (2) | — | 19 | 0.33 | 2 | 0.85 | 18 |
| MNIST test | 10000 (3) | ✓ (30) | 15 | 0.71 | 10 | 0.80 | 12 |
| Olivetti | 400 (2) | ✓ (30) | 38 | 0.73 | 40 | 0.76 | 10 |
| Path based | 300 (2) | — | 2 | 0.68 | 3 | 0.68 | 19 |
| R15 | 600 (2) | — | 16 | 0.98 | 15 | 0.99 | 21 |
| S1 | 5000 (2) | — | 15 | 0.99 | 15 | 0.99 | 23 |
| S2 | 5000 (2) | — | 15 | 0.94 | 15 | 0.94 | 23 |
| S3 | 5000 (2) | — | 15 | 0.74 | 15 | 0.74 | 23 |
| S4 | 5000 (2) | — | 15 | 0.66 | 15 | 0.66 | 23 |
| Seeds | 210 (7) | — | 3 | 0.74 | 3 | 0.74 | 24 |
| Spiral | 312 (2) | — | 3 | 0.79 | 3 | 0.79 | 19 |
| Spirals | 250 (2) | — | 2 | 1.00 | 2 | 1.00 | 4 |
| T4 | 8000(2) | — | 19 | 0.51 | 6 | 0.71 | 26 |
| T5 | 8000(2) | — | 33 | 0.38 | 6 | 0.81 | 26 |
| T7 | 8000(2) | — | 32 | 0.40 | 9 | 0.45 | 26 |
| Zachary | 34 (4) | ✓ (30) | 13 | 0.45 | 2 | 0.65 | 25 |

<sup>1</sup> Number of clusters identified by cutting dendrogram at given criteria. Green: correct clusters were found by analysis. Red: incorrect number of clusters were found by analysis. Yellow: correct number of clusters found but cluster centers do not correspond to correct clusters.
<sup>2</sup> In brackets is the number of dimensions of the data set.
<sup>3</sup> In brackets is perplexity used in t-SNE analysis.
<sup>4</sup> Best cut off value was set by sorting the cluster centers by the highest $\gamma$ ( $\delta^*p$ ) values and choosing the same number of cluster centers that exist on the ground truth.
<sup>5</sup> This study.

**Table S3.** Clusterdp performance at dc 2%.

| Data set name | # points<sup>2</sup> | t-SNE<sup>3</sup> | Automatic |  | Best cut off<sup>4</sup> |  | Ref. |
| --- | --- | --- | --- | --- | --- | --- | --- |
|  |  |  | # clusters | FMI | # clusters | FMI |  |
| 2 level 1 | 504 (2) | — | 9 | 0.98 | 9 | 0.98 | t.s.<sup>5</sup> |
| 2 level 2 | 700 (2) | — | 9 | 0.90 | 9 | 0.90 | t.s.<sup>5</sup> |
| 5 easy | 1000 (2) | — | 5 | 0.99 | 5 | 0.99 | t.s.<sup>5</sup> |
| A1 | 3000 (2) | — | 20 | 0.97 | 20 | 0.97 | 20 |
| A2 | 5000 (2) | — | 35 | 0.97 | 35 | 0.97 | 20 |
| A3 | 7500 (2) | — | 25 | 0.59 | 50 | 0.75 | 20 |
| Aggregation | 788 (2) | — | 11 | 0.74 | 7 | 1.00 | 16 |
| Beeps 20 | 550 (2) | — | 2 | 0.67 | 2 | 0.67 | t.s.<sup>5</sup> |
| Bone marrow | 38 (3) | ✓ (7) | 4 | 0.79 | 3 | 0.71 | 22 |
| Cassini | 1250 (2) | — | 6 | 0.70 | 3 | 0.78 | 4 |
| Cuboid | 250 (3) | — | 16 | 0.54 | 4 | 0.79 | 4 |
| Exclamation 1 | 500 (2) | — | 8 | 0.42 | 2 | 0.67 | t.s.<sup>5</sup> |
| Exclamation 2 | 500 (2) | — | 8 | 0.55 | 2 | 0.63 | t.s.<sup>5</sup> |
| Exclamation 3 | 186 (2) | — | 5 | 0.70 | 3 | 0.69 | t.s.<sup>5</sup> |
| Rodriguez 1 | 4000 (2) | — | 5 | n.d. | 5 | n.d. | 6 |
| Rodriguez 2 | 1000 (2) | — | 7 | n.d. | 5 | n.d. | 6 |
| Flame | 240 (2) | — | 5 | 0.73 | 2 | 0.78 | 17 |
| Jain | 373 (2) | — | 8 | 0.43 | 2 | 0.79 | 18 |
| MNIST test | 10000 (3) | ✓ (30) | 12 | 0.78 | 10 | 0.80 | 12 |
| Olivetti | 400 (2) | ✓ (30) | 34 | 0.69 | 40 | 0.74 | 10 |
| Path based | 300 (2) | — | 2 | 0.68 | 3 | 0.67 | 19 |
| R15 | 600 (2) | — | 15 | 0.99 | 15 | 0.99 | 21 |
| S1 | 5000 (2) | — | 15 | 0.99 | 15 | 0.99 | 23 |
| S2 | 5000 (2) | — | 15 | 0.94 | 15 | 0.94 | 23 |
| S3 | 5000 (2) | — | 17 | 0.75 | 15 | 0.75 | 23 |
| S4 | 5000 (2) | — | 17 | 0.66 | 15 | 0.65 | 23 |
| Seeds | 210 (7) | — | 3 | 0.81 | 3 | 0.81 | 24 |
| Spiral | 312 (2) | — | 3 | 0.92 | 3 | 0.92 | 19 |
| Spirals | 250 (2) | — | 4 | 0.80 | 2 | 1.00 | 4 |
| T4 | 8000(2) | — | 16 | 0.57 | 6 | 0.73 | 26 |
| T5 | 8000(2) | — | 22 | 0.47 | 6 | 0.81 | 26 |
| T7 | 8000(2) | — | 22 | 0.47 | 9 | 0.52 | 26 |
| Zachary | 34 (4) | ✓ (30) | 16 | 0.41 | 2 | 0.94 | 25 |

<sup>1</sup> Number of clusters identified by cutting dendrogram at given criteria. Green: correct clusters were found by analysis. Red: incorrect number of clusters were found by analysis. Yellow: correct number of clusters found but cluster centers do not correspond to correct clusters.
<sup>2</sup> In brackets is the number of dimensions of the data set.
<sup>3</sup> In brackets is perplexity used in t-SNE analysis.
<sup>4</sup> Best cut off value was set by sorting the cluster centers by the highest $\gamma$ ( $\delta^*p$ ) values and choosing the same number of cluster centers that exist on the ground truth.
<sup>5</sup> This study.

**Table S4.** Performance of clustering algorithms on data sets using UMAP to reduce dimensionality.

| Clusterdv performance<sup>1</sup> |  |  |  |  |  |  |  |  |  |  |  |
| --- | --- | --- | --- | --- | --- | --- | --- | --- | --- | --- | --- |
| Data set name | # points<sup>2</sup> | UMAP<sup>3</sup> | Onion |  | Simplex |  | SI |  | Best cut off<sup>4</sup> |  | Ref. |
|  |  |  | # | FMI | # | FMI | # | FMI | # | FMI |  |
| Bone marrow | 38 (2) | (10, 0.1) | 2 | 0.69 | 2 | 0.69 | 3 | 0.82 | 3 | 0.82 | 22 |
| MNIST test | 10000 (2) | (10, 0.0001) | 354 | 0.19 | 280 | 0.22 | 9 | 0.76 | 10 | 0.78 | 12 |
| Olivetti | 400 (2) | (5, 0.5) | 53 | 0.65 | 46 | 0.63 | 46 | 0.63 | 40 | 0.58 | 10 |
| Zachary | 34 (2) | (10, 0.1) | 2 | 0.94 | 1 | 0.71 | 2 | 0.94 | 2 | 0.94 | 25 |
| Clusterdp performance (1% dc)<sup>1</sup> |  |  |  |  |  |  |  |  |  |  |  |
| Data set name | # points<sup>2</sup> | UMAP<sup>3</sup> | Automatic |  | Best cut off<sup>5</sup> |  | Ref. |  |  |  |  |
|  |  |  | # | FMI | # | FMI |  |  |  |  |  |
| Bone marrow | 38 (2) | (10, 0.1) | 13 | 0.41 | 3 | 0.818292 | 22 |  |  |  |  |
| MNIST test | 10000 (2) | (10, 0.0001) | 38 | 0.51 | 10 | 0.722537 | 12 |  |  |  |  |
| Olivetti | 400 (2) | (5, 0.5) | 28 | 0.49 | 40 | 0.579641 | 10 |  |  |  |  |
| Zachary | 34 (2) | (10, 0.1) | 4 | 0.67 | 2 | 0.663605 | 25 |  |  |  |  |
| Clusterdp performance (2% dc)<sup>1</sup> |  |  |  |  |  |  |  |  |  |  |  |
| Data set name | # points<sup>2</sup> | UMAP<sup>3</sup> | Automatic |  | Best cut off<sup>5</sup> |  | Ref. |  |  |  |  |
|  |  |  | # | FMI | # | FMI |  |  |  |  |  |
| Bone marrow | 38 (2) | (10, 0.1) | 18 | 0.361922 | 3 | 0.818292 | 22 |  |  |  |  |
| MNIST test | 10000 (2) | (10, 0.0001) | 19 | 0.680719 | 10 | 0.792613 | 12 |  |  |  |  |
| Olivetti | 400 (2) | (5, 0.5) | 24 | 0.442919 | 40 | 0.573789 | 10 |  |  |  |  |
| Zachary | 34 (2) | (10, 0.1) | 5 | 0.627011 | 2 | 0.663605 | 25 |  |  |  |  |

<sup>1</sup> Number of clusters identified by cutting dendrogram at given criteria. Green: correct clusters were found by analysis. Red: incorrect number of clusters were found by analysis. Yellow: correct number of clusters found but cluster centers do not correspond to correct clusters.
<sup>2</sup> In brackets is the number of dimensions of the data set.
<sup>3</sup> In brackets is the number of neighboring points and minimum distance used in UMAP.
<sup>4</sup> SI dendrogram was cut on the number of clusters of ground truth.
<sup>5</sup> Best cut off value was set by sorting the cluster centers by the highest $\gamma$ ( $\delta^*p$ ) values and choosing the same number of cluster centers that exist on the ground truth.

## Author contributions

JCM and MBO designed the project, developed the clusterdv algorithm, collected data, analyzed data and wrote the manuscript.

## Acknowledgements

JCM was supported by a Scholarship from the Portuguese Fundação para a Ciência e Tecnologia (FCT). The work was supported by grants to MBO from the Bial Foundation (185/12), Marie Curie (FP7-PEOPLE-2011-CIG) and FCT (PTDC/NEU-NMC/1276/2012). We thank Alfonso Renart, Christian Machens and Gonzalo de Polavieja for helpful discussions during the project and Drew Robson, Jennifer Li and Mattia Bergomi for critically reading the manuscript.

## References

1 Jain, A.K. et al. (1999) Data clustering: a review. ACM computing surveys (CSUR)

2 Kleinberg, J. (2002) An impossibility theorem for clustering. NIPS

3 Xu, R. and WunschII, D. (2005) Survey of Clustering Algorithms. IEEE Trans. Neural Netw. 16, 645–678

4 Wiwie, C. et al. (2015) Comparing the performance of biomedical clustering methods. Nat Meth 12, 1033–1038

5 Ester, M. et al. (1996) A density-based algorithm for discovering clusters in large spatial databases with noise. KDD

6 Rodriguez, A. and Laio, A. (2014) Clustering by fast search and find of density peaks. Science 344, 1492–1496

7 Wang, X.F. and Xu, Y. (2015) Fast clustering using adaptive density peak detection. Statistical Methods in Medical Research DOI: 10.1177/0962280215609948

8 Breiman, L. et al. (1977) Variable Kernel Estimates of Multivariate Densities. Technometrics 19, 135–144

9 Sibson, R. (1973) SLINK: an optimally efficient algorithm for the single-link cluster method. The computer journal 16, 30–34

10 Samaria, F.S. and Harter, A.C. (1994) Parameterisation of a stochastic model for human face identification. Proceedings of the Second IEEE Workshop on Applications of Computer Vision

11 Sampat, M.P. et al. (2009) Complex wavelet structural similarity: a new image similarity index. IEEE Trans Image Process 18, 2385–2401

12 LeCun, Y. et al. (1998), Gradient-based learning applied to document recognition. Proceedings of the IEEE.

13 Chen, X. et al. (2016) Infogan: Interpretable representation learning by information maximizing generative adversarial nets. NIPS.

14 Burgess, H.A. and Granato, M. (2007) Sensorimotor Gating in Larval Zebrafish. Journal of Neuroscience 27, 4984–4994

15 Tibshirani, R. et al. (2001) Estimating the number of clusters in a data set via the gap statistic. Journal of the Royal Statistical Society B.

16 Gionis, A. et al. (2007) Clustering aggregation. ACM Trans. Knowl. Discov. Data 1, 4– es

17 Fu, L. and Medico, E. (2007) FLAME, a novel fuzzy clustering method for the analysis of DNA microarray data. BMC Bioinformatics 8, 3

18 Jain, A. and Law, M. (2005) Data clustering: A user’s dilemma. Pattern Recognition and Machine Intelligence

19 Chang, H. and Yeung, D.-Y. (2008) Robust path-based spectral clustering. Pattern Recognition 41, 191–203

20 Karkkainen, I. and Franti, P. (2002) Dynamic local search for clustering with unknown number of clusters. Pattern Recognition.

21 Veenman, C.J. and Reinders, M. (2002) A maximum variance cluster algorithm. IEEE Transactions on Pattern Analysis and Machine Intelligence 24, 1273–1280

22 Monti, S. et al. (2003) Consensus clustering: a resampling-based method for class discovery and visualization of gene expression microarray data. Machine learning 52, 91–113

23 Fränti, P. and Virmajoki, O. (2006) Iterative shrinking method for clustering problems. Pattern Recognition 39, 761–775

24 Charytanowicz, M. et al. (2010) Complete gradient clustering algorithm for features analysis of x-ray images. Information Technologies in Biomedicine 69, 15–24

25 Zachary, W.W. (1977) An information flow model for conflict and fission in small groups. Journal of anthropological research 33, 452–473

26 Karypis, G. et al. (1999) Chameleon: Hierarchical clustering using dynamic modeling. IEEE Computer 32, 68–75

27 Zhang, J. et al. (2016), A robust density-based clustering algorithm for multi-manifold structure. presented at the Proceedings of the 31st Annual ACM Symposium on Applied Computing, pp. 832–838

28 Mehmood, R. et al. (2017) Clustering by fast search and mergeof local density peaks for geneexpression microarray data. Sci Rep DOI: 10.1038/srep45602

29 Macosko, E.Z. et al. (2015) Highly Parallel Genome-wide Expression Profiling of Individual Cells Using Nanoliter Droplets. Cell 161, 1202–1214

30 Dimitriadis, G. et al. T-SNE visualization of large-scale neural recordings. DOI: 10.1101/087395

31 Berman, G.J. et al. (2014) Mapping the stereotyped behaviour of freely moving fruit flies. Journal of The Royal Society Interface 11, 20140672–20140672

32 McInnes, L. and Healy, J. (2018) UMAP: Uniform Manifold Approximation and Projection for Dimension Reduction. Arxiv

33 Martins, S. et al. (2016) Toward an Integrated Zebrafish Health Management Program Supporting Cancer and Neuroscience Research. Zebrafish 13, S–47–S–55

34 Severi, K. et al. (2014) Neural control and modulation of swimming speed in the larval zebrafish. Neuron. 83(3), 692–707.

35 Marques, J.C. et al. (2018) Structure of the Zebrafish Locomotor Repertoire Revealed with Unsupervised Behavioral Clustering. Current Biology DOI: 10.1016/j.cub.2017.12.002

36 Kruskal, J.B. (1956), On the shortest spanning subtree of a graph and the traveling salesman problem. Proceedings of the American Mathematical society, 7, pp. 48–50

37 van der Maaten, L. (2009) Learning a parametric embedding by preserving local structure. Proceedings of the 12th International Conference on Artificial Intelligence and Statistics (AISTATS)

38 Maaten, L. and Hinton, G. (2008) Visualizing data using t-SNE. Journal of Machine Learning Research.

